# Coping with extremes: How Epigenetic and Molecular Adaptations Enable Earthworms to Thrive in Volcanic Soils

**DOI:** 10.1101/2025.07.24.666578

**Authors:** O. Rimington, M. Novo, M.E. Hodson, R. Camarinho, F. Viveiros, C Silva, H. Arruda, A.S. Rodrigues, M. Bruford, S. Short, A.J. Morgan, D. Spurgeon, P. Kille, L. Cunha

## Abstract

Earthworms thriving in naturally occurring geothermal soils offer rare insight into rapid adaptation to environmental extremes. Here, we show that the pantropical earthworm *Amynthas gracilis* survives and flourishes in soils of the Furnas Volcano (São Miguel Island, Azores), where conditions include elevated temperatures (up to 40 °C), high CO_2_ (88.6%), low O_2_ (10%), toxic metals, and mildly acidic pH. In a reciprocal-transplant, mesocosm-based experiment between soils overlying areas of active degassing volcanic gassing (hereafter active degassing soils) and reference soils, convergence of the epidermal thickness of the transplanted earthworms to the resident-soil phenotype (24 ± 3.9 µm active degassing soil, 43.8 ± 8 µm reference soil), was observed within 31 days.

Combining RNA-Seq, DNA (5-cytosine) methylation mapping, and microRNA profiling, this phenotypic change results from coordinated transcriptional and epigenetic reprogramming. While gene-body methylation occurred at ∼98 % of loci, levels varied, and differentially methylated regions were enriched ffor genes with altered expression under volcanic stress. Multi-omics network analysis identified epithelial morphogenesis, circulatory system formation, and neural development as regulatory hubs, highlighted by a set of 41 epithelial-morphogenesis genes showing consistent methylation and miRNA patterns. Additional modules governing ion transport and signal transduction complemented the adaptive response.

Collectively these findings demonstrate that *A. gracilis* employs dynamic DNA methylation and microRNA regulation alongside transcriptional reprogramming to generate a persistent phenotypic adjustment to a volcanic stress. This work advances our understanding of extremophile resilience and provides a scalable model for predicting organismal adaptive capacity in the face of environmental extremes.

## Introduction

Animals and plants living in extreme environments are frequently exposed to a diverse range of stressors, often at varying intensities. To survive, these species evolve integrated suites of anatomical, biochemical, physiological, and behavioural adaptations. Among soil-dwelling invertebrates, characterised by low dispersal and sedentary lifestyles, adaptive plasticity to environmental extremes is especially critical. Earthworms have been shown to adapt to metal pollution (Langdon, et al. 2003; Anderson, et al. 2017; Huang, et al. 2023), high altitude (Perry, et al. 2022), and floodwater inundation (Klok, et al. 2006). However, the case of the ability of the earthworm *Amynthas gracilis* to survive in soils affected by underlying volcanic activity is one particularly striking case (Cunha, et al. 2011; Cunha, et al. 2014), because this species must withstand a combination of high CO_2_ flux (hypercapnia), temperature extremes, low O_2_ (hipoxia), acidic pH, and elevated metal concentrations.

Organisms deploy a suite of conserved biochemical pathways (e.g., heat-shock proteins, metallothioneins, antioxidant enzymes, DNA methylation, and small RNAs) to mount plastic responses to stressors (e.g., (Lindquist and Craig 1988; Morgan, et al. 2007; Zhao, et al. 2016; Wang, et al. 2021). In particular, cell-signalling cascades triggered by oxidative or osmotic imbalance illustrate how rapidly physiology can be reshaped in response to change (Martindale and Holbrook 2002; Wang, et al. 2020). Because these physiological shifts depend on coordinated changes in gene expression, organisms must couple environmental sensing to durable, yet flexible, regulatory systems. Thus beyond the immediate response, epigenetic marks (especially 5-methylcytosine) and non-coding RNAs (notably microRNAs) establish a heritable “regulatory memory” that can mediate non-Mendelian transmission of adaptive phenotypes across sub-Darwinian timescales (Schachtman and Goodger 2008; Geeleher, et al. 2012; Liebers, et al. 2014).

Earthworms are central to global food production as critical ecosystem service providers, modulating nutrient cycling, soil structure, and plant productivity (Fonte, et al. 2023). Understanding the molecular mechanisms underpinning their adaptive plasticity not only enables us to pinpoint species or populations capable of withstanding increasing environmental extremes but also offers a framework for assessing environmental stressor impacts across soil communities. In this study, we therefore focused on defining the underlying gene expression landscapes and two extra-genomic regulators, microRNA abundance and DNA (5-cytosine) methylation, to test their contributions to plasticity in the invasive earthworm *Amynthas gracilis*, following transplantation between soils effected by geothermally “degassing” and reference soils on São Miguel Island.

On the mid-Atlantic island of São Miguel, diffuse degassing affected volcanic soils, such as those in the Furnas area used for this study, are typified by soil temperatures up to 98°C and CO_2_ degassing leading to soil CO_2_ up to 100% leading to potential hypoxia or hypercapnia (Viveiros, et al. 2010; Viveiros, et al. 2015). Furthermore, these soils also contain elevated concentrations of the metals Cu, Pb, and Zn (Cunha, et al. 2011). Given these extreme conditions, it is surprising that soils in the Furnas Volcano area are colonised by three peregrine earthworm species, *Pontoscolex corethrurus* (Family: Rhinodrilidae) from South America, and *Amynthas gracilis and Amynthas corticis* (Family: Megascolecidae) from Asia while native European lumbricids, though widespread elsewhere on the island (Talavera, et al. 2020), are absent from the hydrothermal and degassing zone (Novo, et al. 2015).

The extremophile nature of *A. gracilis* was investigated by examining the suite of traits that enable this species to cope with the volcanism-linked stressors encountered in the Furnas area. It was hypothesised that *A. gracilis* possesses both genetically determined (“constitutive”) traits and traits induced by plasticity. Previously, it was shown that the epidermis of degassing-site *A. gracilis* residents is significantly thinner than that of counterparts from non-degassing regions on São Miguel (Cunha, et al. 2014). This anatomical modification of the gas-permeable surface has been hypothesised to represent a plastic response that reduces diffusion distances and increases gas-exchange efficiency under prevailing anoxic conditions. In the present study, the regulatory molecular mechanisms governing this developmental change and physiological response were investigated. Three tiers of environmental response were quantified using a reciprocal-transplant, mesocosm-based design: (1) the immediate stress response to altered soil conditions, (2) genome-wide transcriptional changes, assessed by RNA-Seq, and (3) the extra-genomic regulatory mechanisms, microRNA abundance and DNA (5-cytosine) methylation, thought to orchestrate this adaptive plasticity. By integrating transcriptomic, methylation patterns, and small-RNA profiling, the reshaping of gene networks and the establishment of heritable regulatory states in *Amynthas gracilis* under geothermal versus reference soil regimes were elucidated.

## Materials and Methods

The Azores archipelago comprises nine volcanic islands in the North Atlantic Ocean, between 36°45’–39°43’N and 24°45’–31°17’W, at the triple junction of the Eurasian, Nubian and North American plates. Due to this geodynamic setting, seismic and volcanic phenomena are common (e.g., Gaspar et al. 2015). São Miguel is the largest (757Lkm^2^) of the nine islands and is located in the eastern part of the archipelago. Across São Miguel, several manifestations of volcanism, including hydrothermal fumarolic fields, cold CO_2_-rich and hot thermal springs, as well as diffuse soil affected degassing sites, are present (Viveiros, et al. 2008; Viveiros, et al. 2010). To assess how earthworms respond to the range of normal to extreme soil conditions present on São Miguel, two field sites, differing in their degassing activity (e.g., with or without thermal and CO_2_ degassing anomalies (with a maximum temperature of 47.3°C and a maximum CO_2_ of 96.5% at 50 cm depth), were selected for a transplant experiment: (a) degassing sites located inside Furnas Volcano caldera (site referred to hereafter as “Furnas”), in the vicinity of the Furnas Village Fumarolic Field, a region that corresponds to an area of anomalous CO_2_ degassing and temperature, and (b) a non-degassing site at the western flank of Fogo Volcano, at Chã da Macela (site referred to hereafter as “Macela”); although located on an active volcano, experience no recognised anomalous degassing or thermal anomaly (Ferreira, et al. 2005; Viveiros, et al. 2023).

Clitellate adult *Amynthas gracilis* were collected from a site in both the volcanic Furnas site (degassing earthworm collection site), 37° 46’ 12’’ N 25° 18’ 14.399’’ W and volcanic non- degassing site Macela (non-degassing earthworm collection site) 37° 45’ 50.76’’ N, 25° 32’ 3.516’’ W by digging and hand-sorting. Collected individuals were randomly assigned to treatment replicates in a reciprocal, factorial transplant experiment, with earthworm origin and soil exposure location as experimental factors (Fig. 1a). Twenty-four hours after collection, earthworms were exposed either to degassing soil (hereafter “V”) from a grass-covered site in the Furnas caldera (37° 46’ 22.8’’ N, 25° 18’ 14.399’’ W), or to non-degassing reference soil (hereafter “M”) from a pasture near the Macela site (37° 45’ 52.02’’ N, 25° 31’ 30.575’’ W). For each treatment replicate, ten individuals were weighed and placed into perforated, cube-shaped mesh bags (12LL volume) filled with local soil. Six replicate mesocosms per origin (Furnas or Macela) were placed at each exposure site (V or M), resulting in four treatments: VV (Furnas-origin in Furnas soil), VM (Furnas-origin in Macela soil), MM (Macela-origin in Macela soil), and MV (Macela-origin in Furnas soil) (Fig. 1b). Mesocosms were installed flush with the soil surface, ensuring continuous contact between soil and mesh for unhindered gas and pore water exchange. They were covered with a waterproof fabric to prevent waterlogging during heavy rainfall and to reduce moisture loss in dry periods. Mesocosms were maintained in situ for 31 days. At the end of the exposure period, mesocosms were removed and earthworms were hand-sorted, survival recorded, and individuals reweighed. Three individuals per mesocosm were flash-frozen in liquid nitrogen for molecular analyses; the remaining specimens were transferred to the laboratory for further processing.

**Fig. 1.**
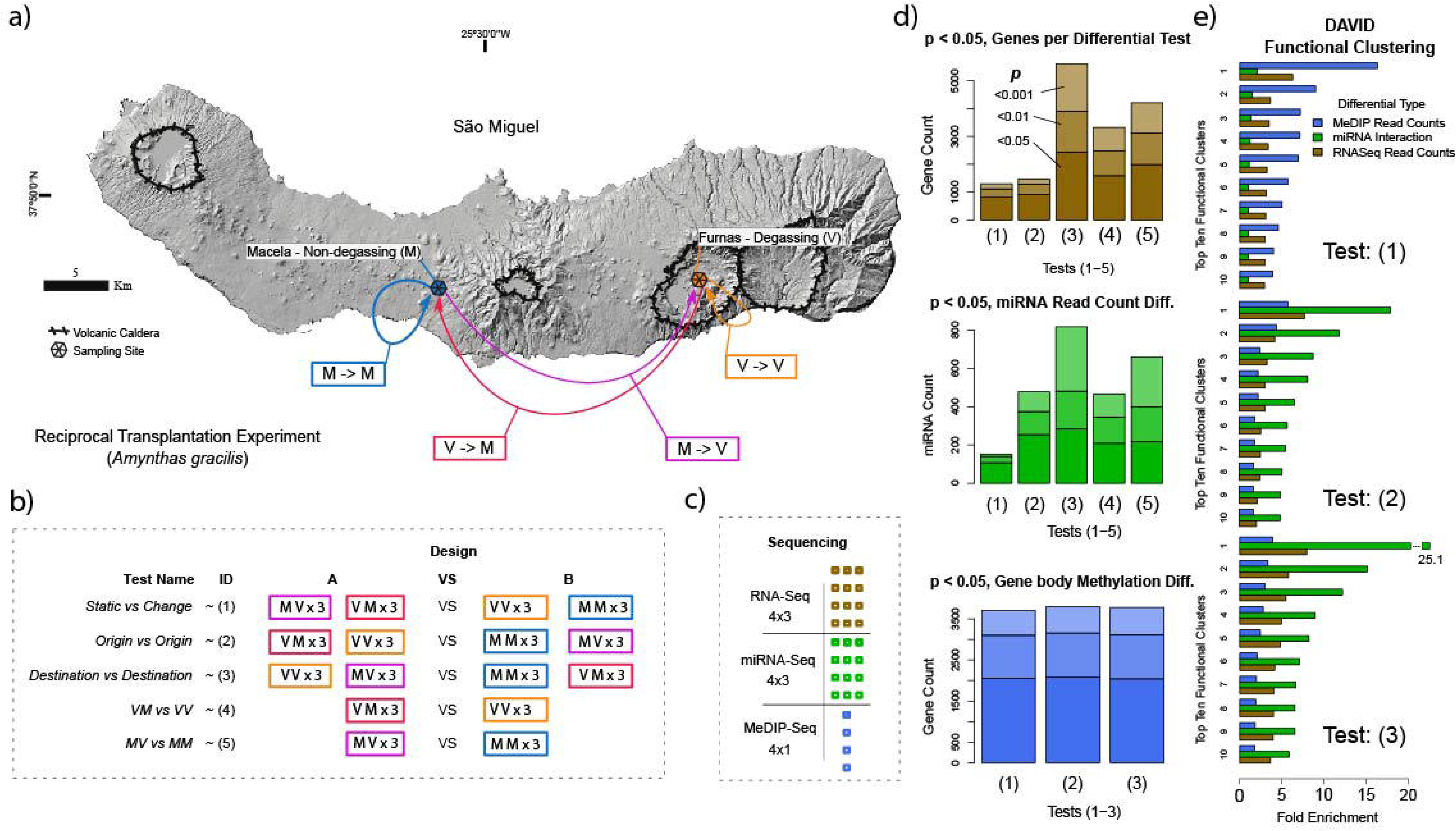
Hill-shaded digital elevation model of São Miguel (Azores) showing the two study areas: Furnas caldera with active soil degassing and elevated temperature (V, orange) and the non-degassing pasture at Macela on Fogo volcano (M, blue). Coloured arrows trace the four transplantation routes used in the 31-day in-situ reciprocal experiment (V → V, V → M, M → V, M → M; three replicate mesh-bag mesocosms per route; 10 adult worms per bag). Black polygons outline volcanic calderas; stars mark the exact sampling/installation sites; scale bar = 5 km. (b) Matrix defining the five statistical contrasts interrogated throughout the study. Each contrast pools replicate libraries from the appropriate source (A) and destination (B) treatments: (1) Static vs Change (resident vs moved worms), (2) Origin vs Origin (degassing vs non-degassing provenance), (3) Destination vs Destination (volcanic vs non-volcanic exposure), (4) VM vs VV, and (5) MV vs MM. Colour coding follows panel a; c) Omics data sets generated: RNA-Seq (4 treatments × 3 replicates), miRNA-Seq (4 × 3), and MeDIP-Seq (4 × 1 pooled replicate). **d)** Numbers of features that changed significantly (Wald test, DESeq2; p < 0.05, 0.01 or 0.001) in each contrast for mRNA genes (brown), miRNAs (green) and gene-body methylated loci (blue). **e)** Top-ten Gene Ontology (GO-BP level 4) clusters from DAVID for the three broad contrasts: (1) *Static vs Change*, (2) *Origin vs Origin*, (3) *Destination vs Destination*. Bars show fold-enrichment contributed by RNA-Seq (dark green), miRNA target interactions (light green) and MeDIP read counts (blue). Prominent clusters include epithelium development, circulatory-system morphogenesis, ion transport, neural development and signal transduction, highlighting convergent pathways underlying rapid epidermal remodelling and physiological acclimation to volcanic stress.

### Environmental characterisation

Soil CO_2_ flux measurements were performed with a portable soil CO_2_ flux station (West Systems S.r.L., Pontedera, Italy) based on the accumulation chamber method (Chiodini, et al. 1998). Soil CO_2_ and O_2_ measurements were performed at the soil surface before starting the experiment following the methodology of Baubron et al. (1991) and gases further sampled at 25 and 50 cm depth using a GA2000 infrared gas detector (Geotechnical Instruments). Even if other volatiles (H_2_S, He, CO, Hg, ^222^Rn) can be released in these environments, CO_2_ represents the main volcanic gas released to these soils (Viveiros, et al. 2010; Silva, et al. 2015; Bagnato, et al. 2018). Soil temperature was measured at the same depths as gas was sampled with a portable T51 Rotronic thermometer. The soil temperature, CO_2_ and O_2_ concentration measurements were performed before the setup and at the end of the experiment inside the mesocosm units.

To characterise soil physicochemical properties, samples were taken from each mesocosm, air-dried, and sieved at 2 mm. The soils were oven-dried for moisture determination at 105 °C. Water-holding capacity was measured following ISO guideline 11274. Organic matter content was measured as loss on ignition at 500 °C in a muffle furnace (Rowell 1994). Soil pH was measured in suspensions of 10 g air-dried soil in 25 ml deionised water following shaking for 15 minutes (ISO 2005). Texture was calculated from the percentage of soil particles in the size ranges < 2 mm, 2 – 63 mm and > 63 mm as determined using a Malvern MasterSizer 200 with a Hydro2000MU wet dispersion unit. A sample of ∼1.5 g of air-dried, sieved soil was analysed with an obscuration of 5 and 25 %. Instrument performance was checked using a Malvern 15- 150 mm quality audit standard and “general purpose sand, 40 – 100 mesh” (Fisher Scientific, Loughborough, UK). Total concentrations of Al, Cr, Cu, Fe, Mn, Ni, Pb, Sr, Ti, and Zn were determined in a 1 g sample of the air-dried soil following *aqua regia* digestion (Arnold, et al. 2008). The certified reference material BCR-143R (Commission of the European Communities, Community Bureau of Reference) was also digested. Digests and method blanks were analysed by inductively coupled plasma-optical emission spectrometry (ICP-OES). Method blanks for Zn were above detection and results were blank corrected, for the other elements method blanks were below detection. Accuracy, as assessed by comparison with the CRM was as follows: Al 127%, Cr 103%, Cu 94%, Fe 104%, Mn 105%, Ni 90%, Pb 110%, Sr 104%, Zn 96%, Ti concentrations were not certified in the CRM.

### Metal body burden by subcellular fractionation

Six earthworms per treatment, one from each replicate for each of the four treatments (total n=24), were selected randomly and depurated for 36h on filter paper before freezing and fractionation (Arnold and Hodson 2007). The frozen earthworms were defrosted, weighed, homogenised in 0.01M Tris-HCl pH 7.5 and fractionated (Arnold et al., 2008) into three different subcellular fractions: a soluble fraction (SOL, comprising the aqueous and cytosolic fraction including soluble proteins such as metallothionein-like and heat-sensitive proteins). an insoluble fraction (MRG comprising metal-rich granules) and a second insoluble fractions (INT, tissue fragments, cell membranes and mitochondria/organelles). Individual fractions were digested in HNO_3_ (Morgan and Morgan 1990), made up to volume with ultrapure water and analysed for the same metals as the soils but by inductively coupled plasma-mass spectrometry, together with As, Bi and Cd. The resulting concentrations are expressed as mg/kg (dry weight) of earthworm tissue.

### Genotyping

All earthworms used were genotyped using the cytochrome oxidase subunit I (COI) gene to confirm their species identity and avoid heterogeneity in the genetic background. Total genomic DNA was extracted from ∼25 mg of tissue from the posterior section or cryogenically powdered tissue. DNAeasy Tissue Kit (Qiagen, Manchester, UK) was used with a final elution with two volumes of 70 µl of TE buffer. A segment of the COI gene was amplified using the primers LCO1490 and HCO2198 (Folmer, et al. 1994). Reactions contained: 4 µl of PCR buffer (Promega, Madison, Wisconsin), 2 µl of 25 mM MgCl_2_ (Promega), 1 µl of 10mM dNTPs, 1 µl of each 10 µM primer, 1 U of Taq DNA polymerase (Promega), 1 µl of genomic DNA template, and sterile H_2_O to a final volume of 20 µl. The PCR profile was 94°C (5 min), 35 cycles of 94°C (30 s), 52°C (30 s), and 72°C (1 min), with a final extension of 10 min at 72°C. PCR products were purified and sequenced by Eurofins (www.eurofinsgenomics.eu) using the same primers. Resulting COI sequences were aligned and compared against reference haplotypes from Novo et al. (2015) to confirm species identity and assess genetic uniformity prior to downstream omics analyses.

### Histological processing and morphometry

Two surviving earthworms from each bag (total n=48) collected after 31 days exposure were depurated for 36 h on damp filter paper to allow them to egest the soil in their gut. The anterior parts (extending three segments posterior to the clitellum) of the purged earthworms were fixed in 4% formaldehyde buffered to pH 7 and stabilized with methanol, dehydrated in a graded ethanol series, cleared in xylene and infiltrated with paraffin wax in a Leica TP1020 (Leica Microsystems, Wetzlar, Germany). The paraffin embedded samples were shaped into histological blocks ensuring the same sample orientation using a Leica Histocentre EG1150 (Leica Microsystems, Wetzlar, Germany). Histological sections (4 μm thickness) were cut on a Leica Rotary microtome RM2035 (Leica Microsystems, Wetzlar, Germany), mounted on albumin-coated slides (Menzel-Glaser, Braunscheig, Germany) and dried at 40°C for 24 h. The slides were then stained with Alcian Blue (Ph 2.5)/ Periodic Acid Schiff (AB/PAS) (Banchroft, et al. 1996). Epidermis thickness of each earthworm was measured in 3 sections (4 fields per section), 40 μm apart. Images were captured using a CoolSNAP-cf camera (Photometrics GmbH, Munich, Germany) coupled to a light microscope Leica DM1000 (Leica Microsystems, Wetzlar, Germany), and analysed with Image Pro-Plus 5.0 (Media Cybernetics, Silver Springs, Maryland). For statistics, the average value from 12 measurements per individual was used for 12 earthworms per treatment. Epidermal thickness measurements were analysed (with or without log_e_ transformation) by two-way ANOVA, with earthworm origin and exposure location as factors, with p≤ 0.05 taken as significant.

### DNA Methylation Immunoprecipitation, RNAseq and miRNA Library Preparation, Sequencing

For Next Generation Sequencing analyses, we selected three replicate mesocosms (bags) out of the six per treatment and used the three previously flash-frozen earthworms from each to be powdered individually (whole body) by pestle and mortar under liquid nitrogen (total n=36). For total RNA extraction, ca. 50 mg of the powder was homogenised in 1.5 ml of Trizol (Life Technologies). After centrifugation at 12,000 g for 5 min, the supernatant was transferred to an Eppendorf and 240 µl of chloroform was added and the mix was incubated for 3 min. The sample was centrifuged at 12,000 g for 15 min (4°C), and the upper phase was mixed with 250 µl absolute ethanol. RNeasy mini columns (Qiagen) were used to clean the RNA, and samples were eluted by centrifugation at 8,000 g in two volumes of 30 µl of H_2_O. Extracted RNA quality, integrity and quantity were checked in a 2100 Bioanalyzer (Agilent) (RNA Nano Chip) and Nanodrop (Thermo Fisher Scientific). Twelve samples (three replicates each comprising pooled RNA from three individuals from the same bag for each of the four transplant conditions) were poly-A selected and prepared for Truseq RNA paired-end cDNA library construction by Edinburgh Genomics (<u>genomics.ed.ac.uk</u>). 100 bp paired-end libraries were multiplexed and sequenced in two Illumina Hiseq 2000 lanes (Illumina Inc).

Small RNAs were extracted from the same samples (36 samples pooled in 12 libraries) with miRNeasy Mini kit (Qiagen Ltd) following the manufactureŕs guidelines. The samples were quality assessed using Bioanalyzer and Nanodrop and supplied to Edinburgh Genomics (<u>genomics.ed.ac.uk</u>), where the libraries were prepared using TruSeq Small RNA library kit (Illumina Inc) and 50 bp single-end sequencing on an Illumina 2500 HiSeq platform using HiSeq V3 chemistry (Illumina Inc).

For the DNA methylation analysis, genomic DNA was extracted from powdered tissue using the same samples as for total RNA and small RNA using the DNeasy kit (Qiagen Ltd). DNA quality was assessed using a nanodrop spectrophotometer (Thermo Fisher Scientific), with integrity and size being determined using a 0.4% agarose gel electrophoresis. For each treatment, equimolar DNA aliquots from three individuals of a single replicate mesh bag, the same as those used in the RNA-Seq workflow, were pooled to ensure sufficient input. Libraries for MeDIP-Seq libraries were constructed following methylated-DNA immunoprecipitation using the DNA Methylation IP Kit (Zymo Research, Orange, CA). Briefly, pooled DNA was sonicated to ∼200–500 bp fragments, denatured, and incubated with 5-methylcytosine–specific antibodies. Immunoprecipitated DNA was recovered, then subjected to a three-step PCR enrichment: first with primers containing partial adapter sequences and four random nucleotides to minimise bias, followed by two successive rounds of amplification to append full-length sequencing adapters and unique barcodes. Libraries were quantified using the Agilent 2200 TapeStation and by qPCR. Sample concentrations were normalised to 4 nM, before sequencing on an Illumina HiSeq 2500 using 100 bp paired-end reads.

### Generating the *A. gracilis* Genome

High-molecular-weight genomic DNA was extracted from a single adult *Amynthas gracilis* collected at the Furnas degassing site using a standard phenol–chloroform protocol. DNA quality was assessed by 1% agarose gel electrophoresis and quantified using a NanoDrop spectrophotometer (Thermo Fisher Scientific). A draft genome was subsequently generated using four paired-end libraries (insert sizes: 350 bp and 650 bp) and two mate-pair libraries (5 kb and 10 kb inserts), sequenced on an Illumina HiSeq 2500 platform. This yielded approximately 100× coverage with 150 bp paired-end reads and an additional ∼10×coverage from mate-pair libraries, with sequencing depth balanced across library types. K-mer analysis (Simao, et al. 2015) indicated a high level of heterozygosity, prompting the use of *Platanus* v1.2.4 for de novo assembly (Kajitani, et al. 2014). The initial graph-based assembly resolved allelic variation by collapsing bubbles representing heterozygous loci. Mate-pair reads were then employed during scaffolding to bridge repetitive regions and connect contigs. Finally, a post-assembly allelic-variant collapse step was performed to reduce redundancy, consolidating residual haplotigs into single consensus scaffolds, resulting in a high-quality haploid representation of the *A. gracilis* genome.

### Sequence Read Mapping Against the Newly Generated Genome

Short-read libraries for all three sequencing experiments were end-clipped and filtered with Trimmomatic (Bolger, et al. 2014). The RNASeq and Me-DIP libraries were aligned to the generated *A. gracilis* genome using BBmap (Bushnell 2014) with the pre-set ‘slow’. RNASeq libraries aligned to the genome at a mean rate of 86.1%, and the MeDIP libraries aligned at 90.5% (see supplementary table S1 for library sizes and alignment rates). SAMtools was used to convert output SAM files to sorted and indexed BAM files (Li, et al. 2009). The ‘htseq-count’ function of the HTSeq python package was used to generate RNAseq data-set read counts. SAMtools (Li, et al. 2009) ‘bedcov’ was used to quantify MeDIP read-count levels within genes and promotor regions. Promotor/TS-factor motif binding levels were divided into two regions: 100 bp 5’ of the transcription start site and 1 Kb 5’ of the transcription start site. These read counts were not used for absolute methylation analysis and did not require normalisation.

Novel miRNA prediction was performed using MiRDeep2 (Friedländer, et al. 2012). Processed reads were aligned to the genome and collapsed into a non-redundant set with the mapper.pl script. The alignments were converted to novel mature miRNA predictions with the miRDeep2.pl script. A bowtie database (Langmead, et al. 2009) was built using a combination of the predicted miRNAs and the latest version of the MiRbase database (Griffiths-Jones, et al. 2006; Griffiths-Jones, et al. 2008). Bowtie was used to map the collapsed reads onto the database using default short read settings, and a custom script was used to extract the raw read counts per miRNA from the output. While our approach diverges from the MirDeep2 processing steps, generating non-normalized miRNA count data was necessary. This ensured a consistent input format for DESeq2, thereby maintaining the uniformity of the statistical pipeline and facilitating reliable downstream comparisons.

Studies in humans and *Drosophila* have discovered canonical cohorts of 1,917 and 258 active miRNAs, respectively, according to miRbase (Kozomara, et al. 2019). The outputs from this processing pipeline included upwards of 20,000 hits per sample. However, the read counts per hit exhibited a power law distribution, with 10,313 miRNA targets achieving an average read count of less than 20. To select the hits most likely to represent active miRNAs, and with reference to typical miRNA cohort sizes, the output set was limited to the top 2,000 hits by mean expression. All the novel miRNAs also fell within this upper range.

### Methylome Gene Models

Custom software called Renoo (https://github.com/OliverCardiff/Renoo) was developed to build methylation gene models for the MeDIP data set. This software reads SAM formatted alignment files and analyses the coverages of genomic annotation elements across normalised spatial intervals. These magnitudes within these intervals are then scaled by a coupling-vector to compensate for CpG density precipitation efficiencies (Wilson and Beck 2016). The coupling vector was defined as the mean observed coverage per sample, divided by the all-sample mean. The tool developed also aggregates the MeDIP read coverage over a set of elementary annotations. For example, the 0-10%, 10-20% … 90-100% intervals along the 5’->3’ length of an exon. The scaled read pileup allows for the profiling of a typical coverage pattern over the set of all spatial element. The software also supports rank- grouping these intervals by an additional variable, in this case, global-average gene expression levels from the corresponding RNA-Seq libraries were used. This process involves sorting the elements into deciles of their parent gene’s all-sample geometric mean expression level and calculating the Me-DIP read coverage intra-group arithmetic means per interval (see Fig. 2a).

**Fig. 2.**
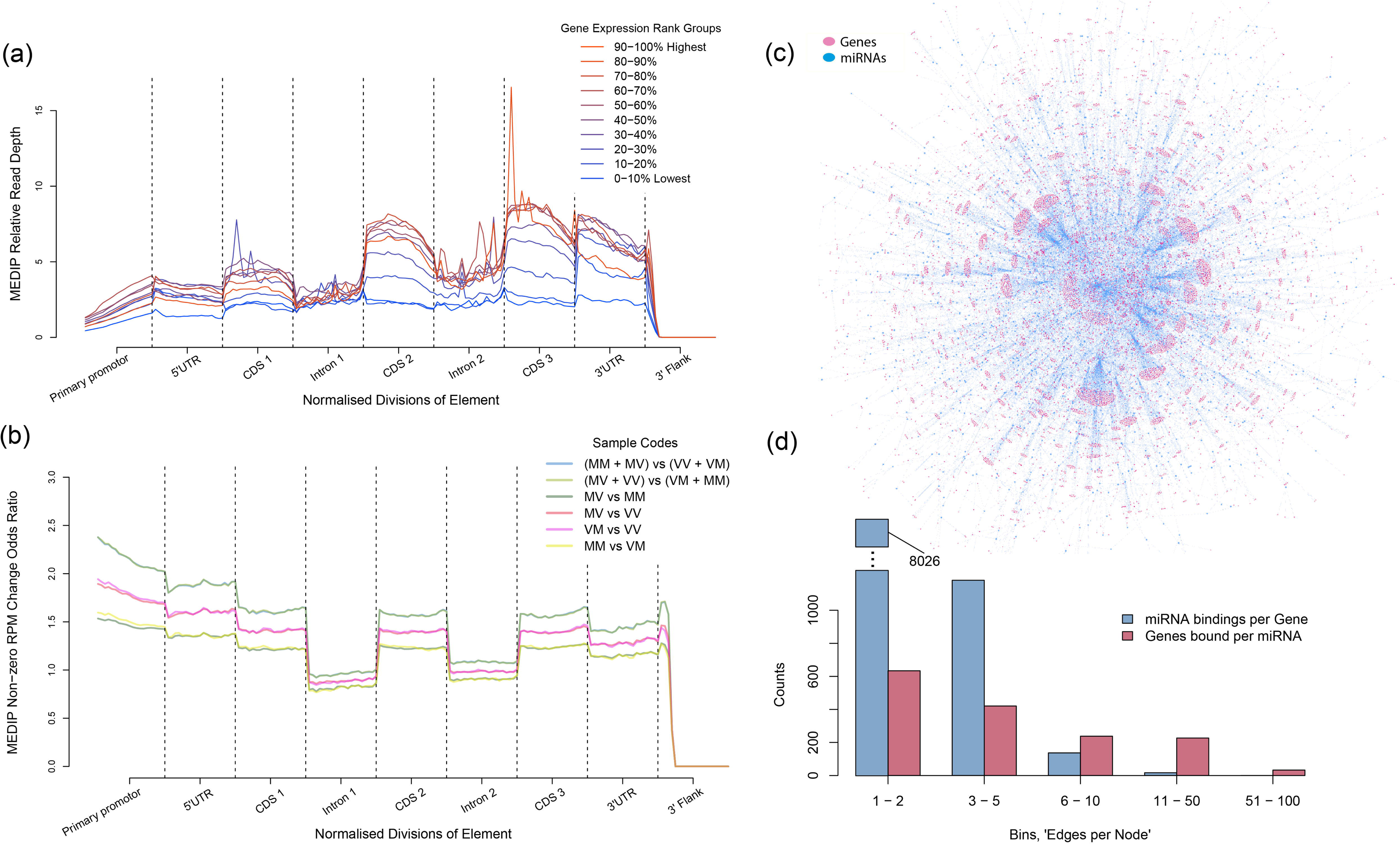
Model gene body methylation spatial representation, (a) Interval-based relative read depth-of-coverage over a model gene, divided into groups corresponding to geometric mean gene expression rank deciles (b) Read-mapping likelihood over a model gene, same data as (a) represented as binary probability of (any) methylation (c) Visualisation of the full miRNA regulatory network between genes (Pink) and miRNAs (blue); (d) Bar graph of the per-node edge count to miRNA to gene interaction relationships.

Because our samples consisted of pooled tissues, where methylation can be highly cell-type–specific (Maegawa, et al. 2010; Lokk, et al. 2014), a binary-coverage model was implemented to estimate the probability of methylation occurrence. In this second pass, window read depths were converted to a presence/absence score, and the frequency of windows with nonzero coverage was reported (e.g., a 0.2 frequency indicates that 20 % of loci in that window exhibited any MeDIP-Seq signal).

To visualize methylation patterns in a unified framework, we constructed “methylation gene-models” composed of the following ordered features: a 34 bp proximal promoter, 5′ UTR, three coding exons (separated by two introns), 3′ UTR, and a 300 bp downstream flanking region. Individual genes lacking one or more annotations were coerced into this template, enabling direct comparison of methylation profiles across heterogeneous gene structures.

### Differential Expression and Methylation Analyses

All differential expression comparisons were conducted using the same DESeq2 pipeline (Love, et al. 2014). The “normal” fold-change shrinkage estimator was applied to maximize stringency and reduce noise from lowly expressed features. This conservative approach was especially important for the more heterogeneous MeDIP-Seq libraries (see supplementary Fig. S1–S3 for count-versus-fold-change and significance plots).

The experimental design (Fig. 1b) defined five key contrasts. For RNA-Seq and miRNA-Seq, three adaptive-response scenarios were tested by comparing two groups of three libraries each: (1) Static vs Change, which evaluates environmental change detection by contrasting earthworms remaining in their native soil (non-degassing soil, MM; degassing soil, VV) against those transplanted to the opposite soil, MV and VM; (2) Origin vs Origin, which probes persistent, origin-specific signatures by comparing all degassing-origin earthworms (regardless of soil, VM, VV) against all non-degassing-origin earthworms (MM, MV); and (3) Destination vs Destination, which assesses acclimation by contrasting all earthworms residing in degassing soil (MV, VV) against those in non-degassing soil (MM, VM). For MeDIP-Seq, the four libraries supported two transplant-specific contrasts: (4) degassing- origin earthworms moved to non-degassing soil (VM) versus degassing earthworms maintained in degassing soil (VV), and (5) non-degassing-origin earthworms moved to degassing soil (MV) versus non-degassing earthworms maintained in non-degassing soil (MM).

To explore miRNA-mediated regulation, the top 2,000 expressed miRNAs were aligned against the set of annotated transcripts, derived via TransDecoder utility scripts in the Trinity package (Haas, et al. 2013), using Bowtie 2 with parameters -k 500 --mp 2 -L 7. This configuration allows up to 500 alignments per miRNA, lowers the mismatch penalty to 2, and reduces the seed length to 7 (the empirically optimal minimum for perfect-match seed mapping of miRNAs; Lewis et al. (2005)), with no soft clipping. No reported alignment exceeded two mismatches. The resulting miRNA–gene network (supplementary Fig. S5 and S6) was then used to interpret primary regulatory impacts in the Destination vs Destination and Origin vs Origin comparisons (the Static vs Change contrast yielded too few significant miRNAs for meaningful network analysis). Finally, we propagated DESeq2’s significant fold-changes for functional enrichment clustering through this network to generate gene lists representing the putative downstream targets of miRNA regulation.

### Functional Enrichment

The functional profiles of differentially expressed genes were assessed using the DAVID web server (Huang, et al. 2007). The genes described in *A. gracilis* are not part of a standardised gene naming ontology, as this earthworm is novel with respect to ‘*omic* analysis. A proteome was translated from a genome-derived transcriptome with Transdecoder (Haas, et al. 2013). This proteome was used to search the Uniprot/Swissprot database with ‘blastp’ (Camacho, et al. 2009). All hits achieving an *e-*value below 5e-05 were retained and used as a symbol translation table for DAVID. The complete list of gene annotations was also provided to the web server as a ‘background’ against which to measure enrichment.

A 3×3 matrix of enrichment analyses was conducted, corresponding to three data types (RNA-Seq, miRNA-Seq, MeDIP-Seq) and three experimental contrasts (“Static vs Change,” “Origin vs Origin,” “Destination vs Destination”; Fig. 1d). For each contrast, gene lists were defined by symbol-translated features exhibiting shrinkage-corrected Wald p < 0.05; for miRNA-Seq, lists were derived by propagating significant miRNAs through the miRNA–gene interaction network to their immediate neighbour transcripts. To ensure consistency and comparability, initial clustering was performed at Gene Ontology Biological Process Level 4, and each resulting cluster was subsequently annotated for Cellular Component via simple enrichment. Fig. 1c shows the omics data sets generated: RNA-Seq (4 treatments × 3 replicates), miRNA-Seq (4 × 3), and MeDIP-Seq (4 × 1 pooled replicate).

For the visualisations generated (Figs. 3–5), overlapping ontology terms were collapsed according to predefined rules: the more specific term was retained unless it overlapped at the same hierarchical depth, in which case the next most significant lower-level term was substituted. This procedure was intended to maximise descriptive resolution while preventing redundant labelling (e.g., “transmembrane ion channel” was not redundantly labelled as both an ion channel and a membrane protein).

**Fig. 3.**
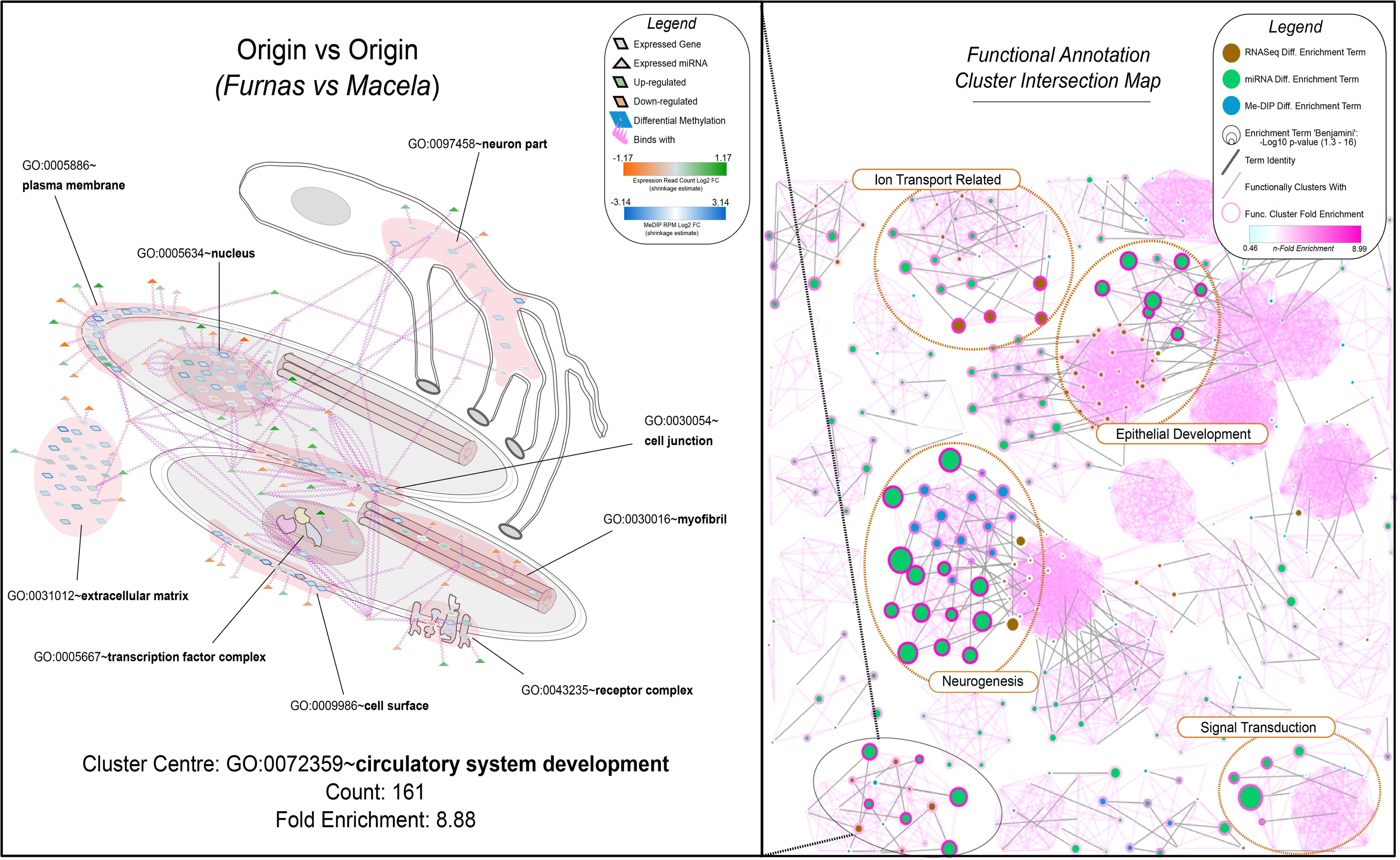
“Origin vs Origin” functional enrichment cluster intersection between data sources (left) and network-view expansion of merged clusters for a single intersection (right). This image shows the miRNA regulatory network for the genes annotation with the term ‘GO:0072359∼circulatory system development’, the network is spatially organised around the GO cellular component terms annotating the cluster, and nodes are coloured by fold-change in the relevant test.

Finally, to produce a unified representation of enriched functions, clusters sharing the same representative GO term across all three contrasts and data types were merged. For example, an “Epithelial Development” trait was defined by the union of all gene lists from clusters containing GO:0060429, with the most significant p-value per cluster being retained.

## Results

### Environmental and Soil Physicochemical Parameters at Degassing and Reference Sites

supplementary table S2 shows the gas and temperature conditions for soils at the degassing (Furnas) and non-degassing (Macela) sites. The mean temperature at a depth of 25 cm measured prior to mesocosm placement was 37 °C (with a minimum of 29.8 and a maximum of 47.3 °C) in the degassing affected transplant site soil, compared to 18°C (with a minimum of 17.8 °C and a maximum of 19.3 °C) for the degassing affected earthworm collection site soil and 17 °C (with a minimum of 16.2 °C and a maximum of 17.6 °C) at the non-degassing site soil. The CO_2_ mean volume was 48.6% (with values between 14% and 96.5%) for the degassing transplantation soil, compared to 6.9% (with a min. of 14% to a max. 96.5%) at the degassing earthworm collecting site and 0.7-0.8% at the non-degassing collection and transplantation sites. The mean O_2_ % (∼10%) was consistent between the degassing transplant site and the degassing earthworm collection site soils. O_2_ levels were higher in the non-degassing site collection and transplant site soils (∼18%), indicating a potential for hypoxia and hypercapnia in soils of the volcanic degassing site.

Soil physical properties indicated differences between the degassing and non-degassing locations, but not between the degassing transplantation and earthworm collection degassing site soils (supplementary table S3). The degassing site soils had a reduced water holding capacity, approximately 50% lower clay content, and a higher sand content than the non-degassing site soils. Degassing soils also had <50% of the organic matter content of the Macela soils (∼11% vs ∼25-35%). Soils at the Furnas degassing site had Pb (108 ±14 mg/kg) and Cu (96 ± 12.1 mg/kg) concentrations ∼6 and ∼5 times higher than those at Macela site soils. Nickel levels at Macela (27.4 mg/kg) were ∼3.5 times higher than in the degassing soils (Table 1). All site soils were mildly acidic (supplementary table S3. All four soil treatments are silt-dominated loams with very low clay content (<5%) and moderate sand (≈35–50%), with lower silt and higher sand content in the V compared to M soil (supplementary table S3).

**Table 1.** Mean concentrations (mg/kg; mean ± SD) of metals measured in soil samples from experimental mesocosms. Conditions represent the four transplant treatments: V–V (degassing-origin earthworms maintained in degassing soil), M–V (non-degassing-origin earthworms exposed to degassing soil), V–M (degassing-origin earthworms exposed to non-degassing soil), and M–M (non-degassing-origin earthworms maintained in non-degassing soil). LoD = Limit of Detection.

| Condition | Al | Cr | Cu | Fe | Mn | Ni | Pb | Sr | Ti | Zn |
| --- | --- | --- | --- | --- | --- | --- | --- | --- | --- | --- |
| LoD | 1.63 | 2.81 | 0.36 | 1.45 | 0.14 | 4.16 | 9.04 | 0.1 | 0.63 | 2.1 |
| V-V | 18600 ± 873.7 | 17 ± 3.6 | 96 ± 12.1 | 19516.7 ± 1662.7 | 587.2 ± 55.4 | 6.2 ± 1.3 | 108.2 ± 14.4 | 34.7 ± 4 | 1074.3 ± 93.5 | 351.8 ± 37.2 |
| M-V | 18633.3 ± 740.9 | 18.1 ± 3 | 100.8 ± 19.9 | 14033.3 ± 6337.9 | 582.8 ± 40.4 | 9.1 ± 5.6 | 108.6 ± 13.5 | 37.3 ± 2.8 | 985.8 ± 123.6 | 385.5 ± 90.4 |
| V-M | 22500 ± 2259.1 | 23.1 ±<br>8.4 | 20 ± 5.1 | 23816.7 ± 4138.2 | 692.8 ± 65.6 | 28.5 ±<br>13.2 | 17.5 ± 4.2 | 73.9 ±<br>30.2 | 3270 ± 770.8 | 155.3 ± 59.6 |
| M-M | 23100 ± 896.3 | 21.5 ±<br>5.4 | 20.2 ± 6.2 | 24366.7 ± 3870.7 | 712.2 ± 51.8 | 26.3 ±<br>12.7 | 17.6 ± 2 | 71.1 ±<br>25.2 | 3266.7 ± 681.4 | 173 ± 79.7 |

### Earthworm Genotyping and Genome Assembly

Mitochondrial COI barcoding of sampled individuals revealed two haplotypes differing by a single nucleotide over 648 bp, consistent with previously described São Miguel *A. gracilis* lineages (Novo, et al. 2015), thereby confirming species identity and genetic uniformity prior to downstream omics analyses.

The generated genome assembly N50 was 478Kb, with 4,350 scaffolds, and a total reconstruction genome size of 589Mb. This genome is available at NCBI Bioproject PRJNA513445. The final gene model identification and annotation was created using MAKER2 (Holt and Yandell 2011). In total, 26,951 gene models were predicted. The genome was 78% complete for Metazoan BUSCO genes (Simao, et al. 2015), with a duplication rate of 11%, a fragmented rate of 4.9%, and a missing rate of 16%. This assembly shows no evidence of allelic inflation and, therefore, likely represents a good haploid representation of the genome. A secondary version of the assembly shows a higher level of completeness indicated through an 87% BUSCO score, but a smaller N50 of 370 Kb. This assembly exhibited a 19% duplication rate, a feature that may indicate that not all loci have collapsed to their haploid counterparts. As such, this genome may contain unphased diploid copies of some genes. For completeness, this version is also available through NCBI Bioproject PRJNA513445.

### Earthworm Tissue Metal Concentrations

Earthworm metal burdens (Table 2) were overwhelmingly partitioned into the internal/tissue fraction (INT), with the soluble pool (SOL) carrying an intermediate load and the metal-rich granule fraction (MRG) the smallest. On average across treatments, INT held roughly 80– 90% of the total body load of each metal, SOL about 5–15%, and MRG only 1–10%. Fe, Cu, Cd, and Pb all adhered to this INT ≫ SOL > MRG hierarchy (Tukey HSD p < 0.05). Granule-bound Al and Mn were significantly enriched compared to their soluble pools (Al: 400 mg/kg vs. 27 mg/kg; Mn: 14 mg/kg vs. 40 mg/kg in MRG and SOL respectively; Tukey HSD p < 0.05), capturing a disproportionate proportion of these potentially reactive elements and thereby limiting their cytosolic availability. For example, Al in INT averaged ∼2 500 mg/kg (VV), approximately 83% of the totalled concentration of Al, versus ∼27 mg/kg in SOL (∼1%) and ∼400 mg/kg in MRG (∼13%).

**Table 2.**
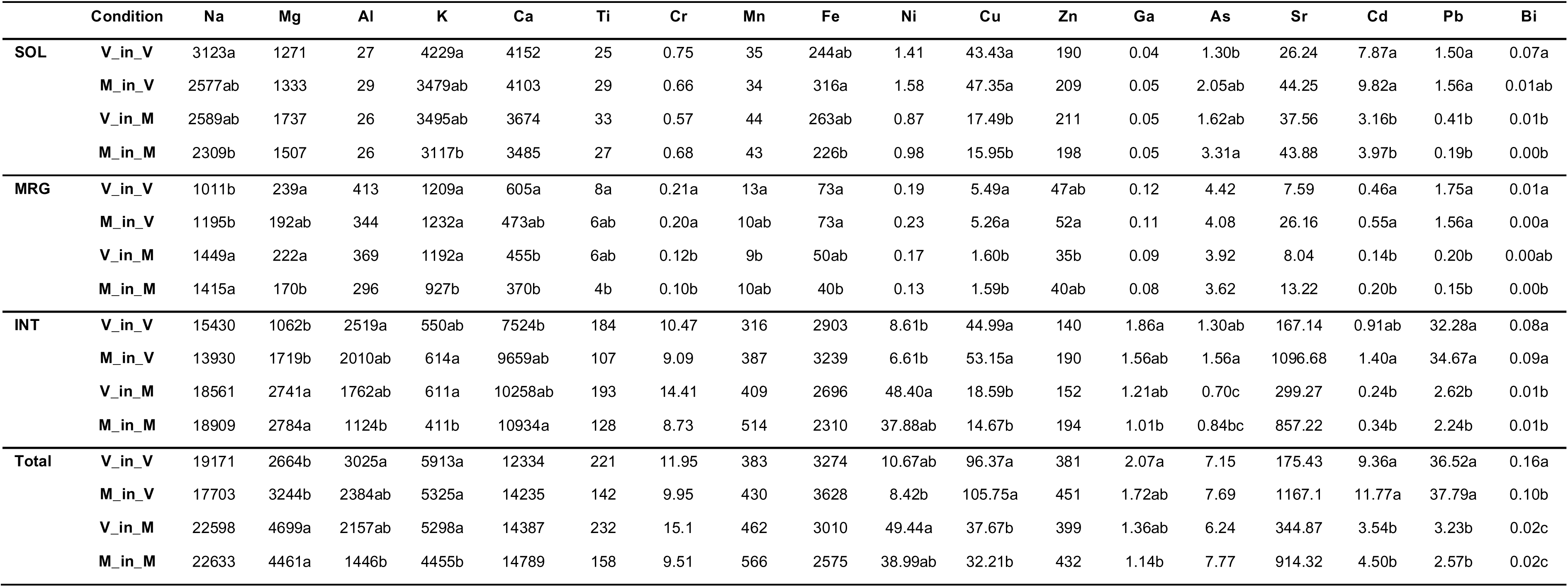
Earthworm body matter metal content (mg/kg) is fractionated between metal-rich granules (MGR), soluble material (SOL), and soft matter (INT: tissue fragments, membranes, etc). Different letters show significant differences after a comparison using Tukey HSD (p < 0.05).

Absolute metal levels were significantly affected by treatment type with the distribution of metals between tissue fractions remaining consistent for each metal. Earthworms in degassing soils (VV, MV) accumulated significantly more total and soluble Cu, Cd, Pb, and Bi than those in non-degassing soils (VM, MM) (all pairwise p < 0.05). Total Cu, for instance, was over three times higher in VV and MV than in MM, and Cd and Pb in SOL increased by 0.5 to 1 times higher under degassing conditions. Conversely, Ni was highest in Macela- exposed worms (VM, MM), with VM showing more than four times the Ni levels of VV (Tukey HSD p < 0.05), reflecting the elevated Ni content of Macela soils.

### Morphometric Changes

After 31 days in either the active degassing (V) or non-degassing (M) soils, earthworm epidermal thickness matched the resident soil conditions regardless of origin (supplementary Fig. S4). In degassing soils, mean epidermal thickness was 24 ± 3.9 μm, whereas in Macela soils it was 42.8 ± 8 μm. Furnas-origin earthworms transplanted to Macela exhibited a 78.3 % increase in epidermal thickness relative to V controls, and M-origin worms moved to V showed a 44.1 % decrease (Tukey HSD, p < 0.05). Body-mass changes of the earthworm showed that those remaining in the active degassing V site soil lost 10.7 % of their initial mass, those remaining in M site soil lost 3.3 %, VM transplantees gained 5.4 %, and MV transplantees lost 13.6 % (supplementary Fig. S4).

### Methylome Models

Of the 26,951 genes analysed, 93.5 – 96.4% were gene-body methylated to some extent (100+ MeDIP reads mapped on average). Fig. 2a shows a visualisation of the consolidated MeDIP read mappings to the common static gene model of methylation read depth for genes. There are clear differences between the rates of read alignment per gene element. The separation in methylation rates occurs most strongly between the lowest five ranks’ groups, indicating that the lower 50% of the gene expression ranks have a positive linear relationship between their epigenetic interactions and their expression level. In contrast the upper 50% of the group showed little correlation. These conclusions are supported by both the relative read depth and methylation probability models.

Overall, coding sequences were more likely to receive highly abundant MeDIP read mapping than promoter regions, with methylation more prominent in the centre of the gene model than towards the 5’ end. However, a binary state comparison of the gene model demonstrates that more individual introns had a higher methylation probability than individual coding sequences (Fig. 2b). This suggests that exonic methylation occurs more abundantly and in more consistent locations, whilst intronic methylation occurs less frequently and in a more diverse set of loci. With some inter-sample variability, 9-12% of primary promoter regions are predicted to be methylated. Methylation probability in the binary model also highlighted that the most regularly methylated intragenic features (at any abundance) were the intronic splice junctions, with an overall rate of 28-34% of the full set of junctions methylated to some degree across the four samples in the gene model. This suggests these splice-junctions are the gene elements that most broadly interact with DNA methylation.

### miRNA Networks

An initial miRNA network was generated based on the set of identified miRNA-to-gene binding relationships. In total, 168 novel mature miRNAs were predicted. This network visualisation and edge-distribution summary are shown in Fig. 2c and 2d. Of the 26,951 genes, 9,363 (34%) had at least one miRNA binding site on the conservative rule of two base changes or less compared to the mature sequences. Of these, 8,026 (85.7%) genes were bound by one miRNA, with the rest binding multiple (Fig. 2d). For the 2,000 putative miRNAs included in the alignment query, 1,554 (77%) were indicated to bind to one or more genes, suggesting some degree of over-inclusion error given the arbitrary selection cut-off. Of these miRNAs, 623 (37.4%) only bound a single gene, the rest binding multiple genes, including a small number of highly connected genes (Fig. 2d). Since miRNA functional impact was assessed relative to the bound genes, unbound miRNAs were not included in the annotation enrichment group analysis.

For functional analysis, the abundances of p-significant genes related to miRNA changes in this, and later sections are gathered based on the p-significance of the binding miRNAs. Thus, the functional description of the miRNA networks involved in expression change between samples was performed based on gene, rather than miRNA, annotation. The obtained differential miRNA expression sub-networks are shown in supplementary Fig. S5 and S6 for the “Origin vs Origin” and “Destination vs Destination” respectively, as the “Static vs Change” test did not yield enough p-significant results to generate a usable network. In both possible experimental comparisons, functional enrichment clustering of genes annotated by Gene Ontology BP4 in DAVID yielded similar sets of results to the clustering experiments for the other analyses. Both networks included a highly enriched proportion of membrane-bound proteins. Neural system development was featured in multiple highly abundant clusters in the “Origin vs Origin” test (Counts: 130, 46, 45, 11) and in two less abundant but still highly significant clusters in the “Destination vs Destination” test (Counts: 31, 19). The “Destination vs Destination” network also showed a cluster of 27 ion channel-related genes and a significant gene cluster for vasculature development (Count: 20) and blood circulation (Count: 16). A cluster of 41 genes within the category “morphogenesis of an epithelium” was significant for “Origin vs Origin” test (supplementary Fig. S5).

### Expression Pattern Assessment

The most substantive expression pattern difference between sample groups was in the “Destination vs Destination” test (Fig. 1d). In this comparison, both the miRNAseq and RNAseq datasets showed a pattern of significant p-value counts differential with each treatment comparison (Fig. 1d and 1d). The greatest difference between the number of significant features between the miRNA and RNAseq datasets was for the “Origin vs Origin” case. For this comparison, a stronger signature of effects was evident for the miRNA dataset (middle panel Fig. 1d) compared to the RNAseq dataset (top panel Fig. 1d).

Methylation sample variance was consistently bigger than for the other two ‘omics datasets, likely linked to the lack of replication, use of whole-body tissue-derived DNA and potential method variations (supplementary Fig. S3). Despite this, a set of consistent changes in gene-body methylation in the experimental comparisons was found that showed an identifiable functional specificity as seen also for the RNAseq and miRNA datasets. Overall, the number of *p*-significant changes in the MeDIP data was broadly similar (∼3,000) for the three comparisons, although the significant gene sets differed markedly. In the “Static vs Change” MeDIP comparison, the significant gene set generated functional cluster fold enrichment scores higher than for any other gene list (top panel Fig. 1e), including the other two methylation tests, as displayed by the DAVID Functional Clustering (Middle and Bottom panel Fig. 1e).

### Functional analysis of changes in the RNA and DNA methylation datasets

Enrichment cluster intersection maps in the right-hand portions of Fig. 3-5 show the most significant cluster terms per data source (RNAseq, miRNA, MeDIP). This network view enables comparison of enrichment effects on identical terms across data sources. Notably, every top term identified for a given data type overlaps with at least one enrichment cluster from another source (intersections labelled using the lowest *p*-value term). Furthermore, terms and clusters unique to a single data source are among the least enriched or significant.

Comparing between Fig. 3-5, the same intersections are identified, however enrichments and significances vary by data source. The top emergent cluster intersections are shown in Fig. 6 as a trait matrix. These merged clusters summarise the detailed patterns of effects within five main identified functional categories: 1. Circulatory system Development (GO: 0072359); 2. Epithelial Development (GO: 0060429); 3. Ion Transport (GO:0006811); 4. Neuron Development (GO: 0048666); 5. Signal Transduction (GO: 0007165). The net up/down regulation of genes annotated with these terms along with the associated miRNA expression changes are also shown in the trait matrix. There is a consistent pattern by which more genes with significantly upregulated miRNAs were found in the first four functional categories for the “Destination vs Destination” test (lefthand column plots Fig. 6), whereas the same categories in the “Origin vs Origin” test show a net down-regulated miRNA-gene component (Middle column plots Fig. 6).

Reflecting the functional enrichment differences in the MeDIP-Seq tests, significant differential methylation was most common in the “Static vs Change” test for the functional categories: ‘ion transport’, ‘neuron development’ and ‘signal transduction’. In this test, more genes were affected than in any other data source comparison (compare righthand column plots to lefthand and middle Fig. 6). Methylation significant change rates for the functional categories of ‘circulatory system development’ and ‘epithelial development’ were consistent across the comparisons, suggesting that these levels may be more reflective of a conserved function change identified out with the inherent noise in the dataset.

## Discussion

Our multi-tiered analysis reveals how *Amynthas gracilis* orchestrates morphological, physiological, and molecular responses to thrive in volcanic environments. Reciprocal transplants between degassing V soils and reference M soils uncovered consistent patterns of metal handling, tissue remodelling, and multi-omics reprogramming that underpin this earthworm’s remarkable plasticity.

### The *Amynthas* Genome System and Its Contributions to Function

The assembled genome of the megascolecid earthworm *Amynthas gracilis* enables an unprecedented integrative analysis of its adaptive responses to extreme volcanic-soil conditions. Until now, studies of soil-invertebrate adaptation have primarily focused on either transcriptional changes alone (Bhambri, et al. 2018; Novo, et al. 2020) or on coarse, genome-wide assessments of DNA-methylation shifts (Srut, et al. 2017; Newbold, et al. 2019). By contrast, our work combines physiological, transcriptomic, miRNA, and DNA-methylation data to generate a holistic view of phenotypic plasticity and regulatory network remodelling under geothermal stress.

Consistent with prior invertebrate studies showing that methylation predominantly marks gene bodies and transposable elements (Mandrioli 2007; Suzuki, et al. 2007; Sarda, et al. 2012) (Guynes, et al. 2024), we find that 95–98% of *A. gracilis* genes bear some level of methylation. However, our differential-methylation models reveal that promoter regions, although less heavily methylated overall, undergo disproportionately large methylation changes in response to environmental shifts. This suggests that in earthworms, as in other invertebrates, promoter methylation may act as a dynamic regulatory switch, even if it constitutes a minority of total marks. Up to now, divergence from vertebrates has been attributed to the absence of promoter methylation and higher levels of gene body methylation in invertebrates (Keller, et al. 2016).

Intronic splice-junctions exhibited particularly high methylation probabilities (28–34% of all junctions; Fig. 2b), hinting at a conserved, epigenetic role in alternative-splicing regulation. In vertebrates, methylation near exon–intron boundaries can modulate RNA-pol II kinetics and splicing-factor recruitment, e.g. by preventing CTCF binding and altering Pol II pausing to favour exon skipping (Shukla, et al. 2011) or by recruiting methyl-CpG–binding proteins that guide splicing regulators (Du, et al. 2015). Although our current RNA-Seq analysis addressed gene-level expression, future isoform-resolved studies (e.g. MeDIP-Seq combined with long-read RNA-Seq) could directly test whether junction methylation fine-tunes transcript diversity under volcanic stress, a heritable “splicing code” facilitating rapid adaptation.

Substantial inter-individual variability in gene-body methylation was observed across transplant treatments, mirroring the concept of variable methylation regions (VMRs) as facilitators of heritable phenotypic diversity around a mean trait value (Feinberg and Irizarry 2010). Functional clustering of sites showing differential methylation, gene expression, and miRNA regulation consistently highlighted developmental and signalling pathways (e.g., epithelial morphogenesis, circulatory-system development, neuronal differentiation, ion transport, and signal transduction). This concordance suggests that stochastic methylation changes not only introduce heritable phenotypic variation, but also selectively target conserved adaptive pathways, perhaps reflecting convergent epigenetic strategies across distant taxa (Alakarppa, et al. 2018; Klupczynska and Ratajczak 2021).

Individual miRNAs were found to bind dozens of targets, paralleling the extensive regulatory reach noted in mammals (Friedman, et al. 2009). A pronounced “origin effect” — defined as persistent molecular differences attributable to native-soil history — was observed in miRNA profiles, exceeding that seen in mRNA expression. In earthworms from degassing soils, a net reduction in miRNA abundance was recorded, which is expected to diminish post-transcriptional repression of adaptive genes and, thus, maintain an epigenetic “memory” of prior environmental exposure, potentially bolstering resilience under variable conditions.

The global action of miRNAs has been characterised as predominantly repressive (Cai, et al. 2009; Bartel 2018). This repressive trend was recapitulated in our analyses of the Epithelial Development and Circulatory System Development pathways, where inverse correlations between net miRNA and net mRNA regulation were detected in both “Destination vs. Destination” and “Origin vs. Origin” comparisons. Specifically, miRNA levels were found to be upregulated in the “Destination vs. Destination” test and downregulated in the “Origin vs. Origin” test, implying that, during acute acclimatisation, miRNAs act chiefly to suppress their transcriptional targets. Conversely, the reduction of miRNA abundance in degassing-origin earthworms suggests that repression of adaptive trait–associated transcripts is selectively relieved, thereby preserving expressed phenotypes that may confer adaptive advantage in fluctuating environments.

### Physiological acclimation and adaptation

Pronounced epidermal remodelling was induced by soil transplantation, independent of earthworm origin. In previous work, earthworms maintained in degassing Furnas soils exhibited a thinned epidermis (∼24 µm), whereas those in non-degassing Macela soils retained the resident-soil norm (∼40 µm)(Cunha, et al. 2011). Such adjustments parallel morphotype changes documented in hydrothermal-vent polychaetes (Andersen, et al. 2006) and are consistent with an adaptive response to high CO_2_, low O_2_, and elevated temperature. Transcriptomic analysis identified 181 genes in the epithelial-development pathway, with a >5 times enrichment, confirming the breadth and strength of the genetic program driving this morphological shift. Notably, this pathway was activated in both native and transplanted individuals, indicating true acclimatisation rather than tissue damage. Conversely, epidermal restructuring and cellular alterations in oligochaetes has been related to metal or drought stress and as a protection against desiccation and microbial invasion, demonstrating that abiotic challenges can bidirectionally modulate cuticle architecture (Cunha, et al. 2011; Wang, et al. 2011; Briones and Álvarez-Otero 2018; Kumari, et al. 2024). Body-mass trajectories mirrored these barrier changes: VM transplants gained weight, suggesting metabolic relief (Daniel, et al. 1996), whereas MV transplant worms lost most weight, reflecting the higher physiological toll of survival in active degassing soils (supplementary Fig. S4). The rapid convergence of epidermal thickness on destination values underscores its utility as a sensitive, short-term bioindicator organism stress.

Epigenetic profiling and miRNA data further supported the centrality of epithelial restructuring as a response to volcanic extremes. In the “Static vs Change” comparison, transcript and miRNA pathway-level responses were neutral overall (Fig. 6), yet differential methylation within this same contrast yielded a tightly focused network of epithelial-morphogenesis genes (Fig. 3–5). This suggests that DNA methylation plays a targeted role in regulating barrier pathways. Comparable patterns in crayfish link gene-body methylation to expression stabilisation via chromatin remodelling (Jeltsch and Jurkowska 2014; Gatzmann, et al. 2018), hinting that similar mechanisms may control transcription-factor access during epidermal adaptation in *A. gracilis*.

Although earthworms possess both a closed, haemoglobin-based vasculature for gas transport and an open, hydrostatic vascular network for locomotion (Rieger and Purschke 2005; Reiber and McGaw 2009; Monahan-Earley, et al. 2013), GO annotations for circulatory system development may reflect shared evolutionary origins rather than discrete functional modules. Crucially, however, the epidermis itself plays a central role in cutaneous gas diffusion (Mendes and Nonato 1957), suggesting that genes driving vascular morphogenesis in our datasets may actually contribute to enhancing oxygen uptake and CO_2_ release through the skin.

Merged-cluster analyses, differential expression, and DNA-methylation changes all implicate both angiogenic and morphogenic remodelling of the integument (Fig. 4 and 6). Notably, the number of RNA-seq significant genes in circulatory pathways differs markedly between “Origin vs Origin” and “Destination vs Destination” contrasts, indicating that vascular restructuring is enacted as an active acclimation response rather than a fixed developmental trait. The “Static vs Change” test further reveals stochastic methylation variation on these same genes, highlighting DNA methylation’s role in fine-tuning circulatory remodelling under each O_2_/CO_2_ regime.

**Fig. 4.**
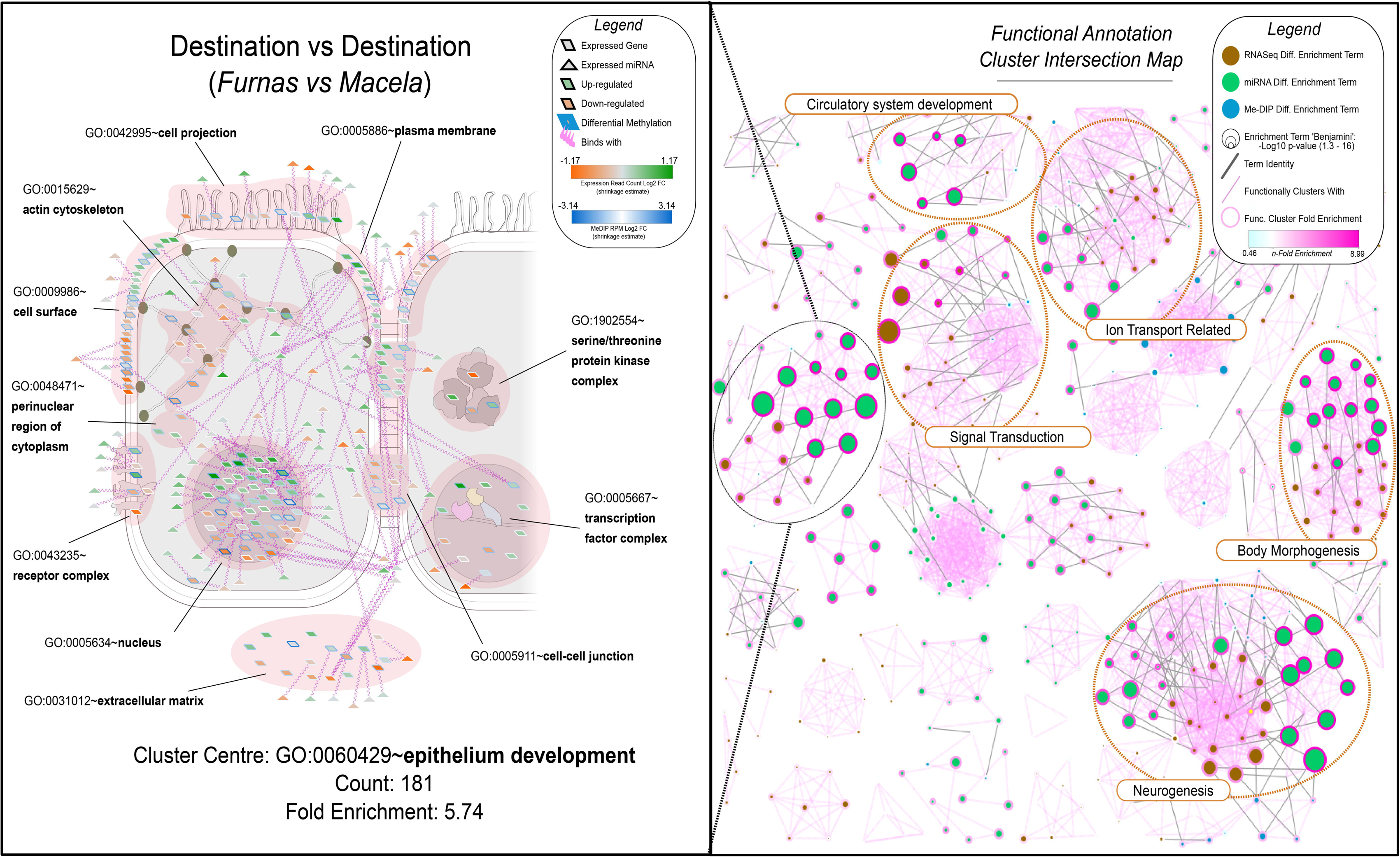
“Destination vs Destination” functional enrichment cluster intersection between data sources (left) and network-view expansion of merged clusters for a single intersection (right). This image shows the miRNA regulatory network for the genes annotated with the term ‘GO:0060429∼epithelium development’, the network is spatially organised around the GO cellular component terms annotating the cluster, and node are coloured by fold-change in the relevant test.

Neuron development was the third morphogenic trait category highly enriched in all three comparisons, albeit in different ways. Changes in neural development can be viewed in conjunction with changes in the signal transduction annotation cluster, as the two overlap considerably via their constituent genes. Signal transduction is a very broad category and can be difficult to disaggregate. However, there was a clear pattern for neurogenesis-associated terms to comprise a substantial part of this subset. It has been shown that invertebrates such as *Drosophila melanogaster* possess dedicated O_2_ and CO_2_ neural or olfactory chemical signalling pathways (Luo, et al. 2009). Our results suggest that similar pathways may be triggering the morphometric changes seen in the epidermis and associated circulatory system. Further, hypoxia encountered by the volcanic site earthworms has been shown to substantially affect neural stem cell differentiation. Research has shown neuronal migration defects and axon pathfinding changes in *Caenorhabditis elegans* (Chang and Bargmann 2008), reducing ion concentrations and leading to hyper-polarization in *D. melanogaster* (Gu and Haddad, 1999) and cation co-transporter activity reduction and resultant hyper-polarization in *Lymnaea stagnalis* (Silverman-Gavrila, et al. 2009).

Most of the literature concerning neural responses to hypoxia in invertebrates concerns membrane-based changes, with little consideration of differentiation and growth alterations (Mannello, et al. 2011). This is reflected in Fig. 5 where the abundance of membrane-bound proteins in the miRNA-regulation network is displayed. A substantial number of neurogenic genes are also identified here, suggesting a role as a more general plasticity response. Signal transduction pathways were generally strongly differentially methylated in the “Static vs Change” test compared to any other comparisons. Neural development-specific pathways also nearly double the rate of differential methylation in this test. This suggests that the ‘general’ environmental change response exhibited may constitute the epigenetic modification of many of the same signalling pathways regardless of the specific nature of the transition encountered by the earthworms. Uniquely, the signal transduction merged cluster also showed more RNAseq differential significance in the “Static vs Change” test, indicating a more general change response.

**Fig. 5.**
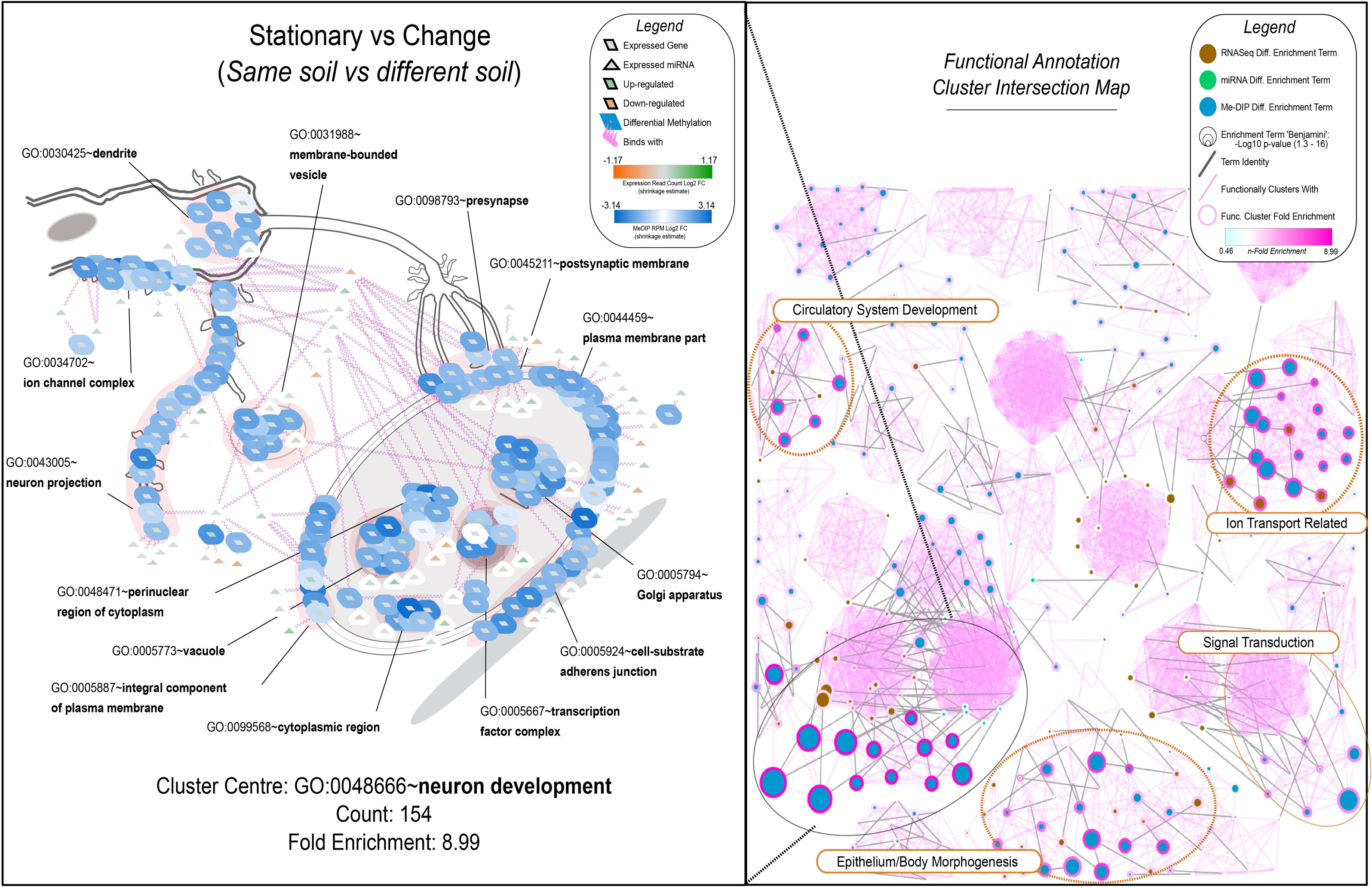
“Static vs Change” functional enrichment cluster intersection between data sources (left) and network-view expansion of merged clusters for a single intersection (right). This image shows the miRNA regulatory network for the genes annotated with the term ‘GO:0060429∼epithelium development’, the network is spatially organised around the GO cellular component terms annotating the cluster, and nodes are coloured by fold-change in the relevant test.

**Fig. 6.**
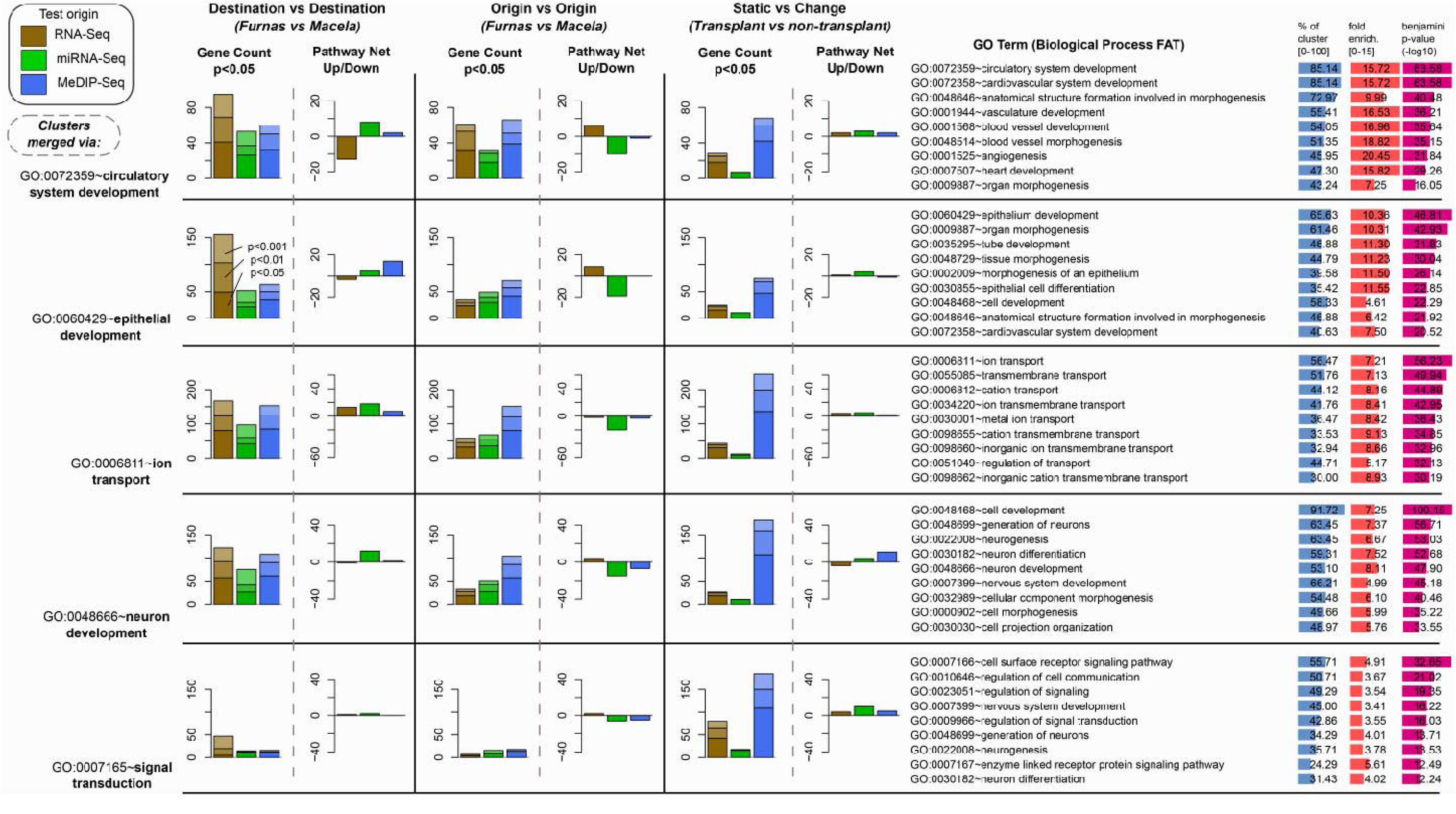
Expression changes for the “Destination vs Destination” (left-hand column charts), “Origin vs Origin” (Middle column charts) and “Static vs change” (right-hand column charts). Data represent the MeDIP, RNAseq and miRNA datasets and select for the largest functional annotation cluster intersections from Figures 3-5. For each comparison (major column), the left-hand sub-column displays the number of genes showing statistically significant changes within each functional annotation cluster. In contrast, the right-hand sub-column illustrates the net up- or down-regulation for each cluster by data source. Additionally, the bar charts on the far right highlight the top nine terms in each cluster, ranked by their enrichment p-values.

### Metal Homeostasis via Compartmentalization and Molecular Drivers of Metal Detoxification

Across all treatments, earthworms partitioned metals consistently into three subcellular pools, INT ≫ SOL > MRG, revealing an evolutionarily conserved sequestration strategy. The INT fraction, encompassing tissue fragments, membranes, and organelles, likely reflects both the incorporation of essential metals into metabolic pathways and the non-specific adsorption of excess metals onto cellular structures. The SOL fraction acts as a dynamic buffer, modulating the immediate bioavailability of metals via soluble proteins, such as metallothioneins, and other cytosolic constituents (Höckner, et al. 2015; Swart, et al. 2022). Finally, the MRG pool provides a dedicated detoxification sink, especially prominent for Al and Mn, thereby shielding critical biomolecules from redox-driven or catalytic damage.

Treatment-specific shifts in metal burdens mirrored soil geochemistry without altering this hierarchy. In V soil exposed worms, soils enriched in Cu, Cd, Pb, and Bi drove elevated accumulation in both INT and SOL fractions. In contrast, earthworm kept in M soil showed pronounced Ni uptake, predominantly allocated to the INT compartment, transplants, reflecting site geochemistry. For As concentration were elevated especially for the MV treatment indicating that the transplanted earthworms failed to regulate internal concentration through mechanisms such as metallothionein binding or granular sequestration to the same extent site as native earthworms in the VV treatment. This observation in consistent with the known toxicity of As and prior reports of epigenetic disruption playing a role in these effects (Kille, et al. 2013). Indeed, that the methylation differentials for ion-transporter genes were markedly higher in the “Static vs Change” comparison, suggesting a broad-scale epigenetic response to prevailing exposure conditions linked to changes in metal trafficking requirements.

The distribution of metals across different cellular fractions suggests that intrinsic sequestration capacities, acting consistently at different exposure levels, govern earthworm resilience to metals. The conserved INT >> SOL > MRG partitioning defines a robust detoxification framework adaptable to diverse metal loads associated with soils impacted by volcanic degassing conditions and anthropogenic contamination. Key pathways involved in this response can be understood from our combined gene expression and epigenetic analyses

Differential omics analyses, including transcript expression, miRNA targeting, and DNA methylation mapping, identified an ion-transport annotation cluster of over 200 genes with significantly altered regulation. Notably, 36.4 percent of these genes encode metal-ion transporters, central to cellular metal homeostasis. Metallothioneins, small cysteine-rich proteins synthesized in the Golgi and likely localized within the SOL fraction, emerged as principal metal detoxification agents (Höckner, et al. 2015). Consistent with their established roles in Zn and Cd binding, SOL abundances of these metals in earthworms exposed to degassing soils suggest metallothionein-mediated regulation, whereas Cu regulation via this mechanism is more nuanced in its response (Swart, et al. 2022).

Beyond Zn and Cd, Pb exhibited increased storage in the INT fraction of active degassing V soil exposed earthworms. For Pb this increase was especially noticeable, with this metal likely bound to phosphate-ligated granules in chloragogenous tissue in this fraction (Morgan and Morgan 1990). This site effect on metal levels is mirrored by a consistent set of miRNA interactions in both “Destination vs Destination” and “Origin vs Origin” comparisons, indicating coordinated post-transcriptional control.

While epidermal thinning under low-O_2_, high-CO_2_ conditions appears to be a general plastic response among oligochaetes, *A. gracilis* ability to tolerate and even thrive in extremely elevated metal environments may define its success in V soils. Our integrated analyses show that *A. gracilis* combines constitutive expansions of detoxification gene families with highly plastic regulatory control of metal-handling pathways. Comparative genomic analyses of ion-transport and detoxification repertoires across co-occurring lumbricid and megascolecid species would clarify whether *A. gracilis* possesses uniquely enriched or more rapidly inducible metal-handling loci, and whether metal tolerance, rather than hypoxia alone, underpins its success in volcanic soils.

### Heat Exposure and Hypoxia-Driven Responses

Individual genes, particularly those encoding heat-shock proteins, chaperonins and other stress-induced factors, are well known to respond transcriptionally to elevated temperatures (e.g. Schoville, et al. 2012; Lockwood, et al. 2017). Similarly, the trehalose transporter Tret1- 2, which mediates the accumulation of the relatively heat-stable disaccharide trehalose, is upregulated under combined temperature and drought stress (Lopienska-Biernat, et al. 2019; Perez and Aron 2023). However, among the main group of annotation clusters, GO Terms linked to exposure to heat stress (e.g. GO:0031072: Heat shock protein binding; GO:0009408: Response to heat; GO:0034605: Cellular response to heat) were, perhaps surprisingly, not among the most enriched sets of pathways identified.

Despite extreme soil temperatures, >40°C in degassing sites, indicating potentially significant heat exposure that exceeds the thermal tolerance of many temperate earthworm species (Viljoen and Reinecke 1992; Edwards and Bohlen 1996). However, despite this evidence heat tolerance, *Amynthas gracilis* showed neither a canonical heat-shock response nor enrichment of GO terms such as GO:0031072 “Heat-shock protein binding,” GO:0009408 “Response to heat,” or GO:0034605 “Cellular response to heat.” It should be noted, however, that our reliance on GO annotations derived from non-annelid models may obscure uncharacterised earthworm heat-stress genes. Native to South-East Asia, where soils often exceed 30 °C, *A. gracilis* nevertheless survives temperatures above the cocoon- viability threshold of related species (Johnston and Herrick 2019). Instead of heat-stress pathways, our data highlight transcriptional, epigenetic, and miRNA signatures linked to hypoxia, pathways governing circulatory and neural development, and epidermal remodelling, suggesting that O_2_ depletion and high CO_2_, rather than heat per se, drive its adaptive response in volcanic soils.

## Conclusion

*Amynthas gracilis* earthworms were reciprocally transplanted between non-degassing and degassing soils, the latter having elevated temperatures, CO_2_ emissions, O_2_ depletion, and altered metal concentration profiles including higher Cu, Pb and Zn. RNAseq, miRNAseq, and MeDIP-Seq profiles were generated to assess functional changes in the response of these earthworms to extreme conditions. Three main differential tests were used to assess adaptive and general environmental responses. *A. gracilis* was found to have highly methylated gene-bodies, with variable gene-component rates, and a clear relationship between mean expression levels and methylation. Independent functional enrichments of significant gene-based differences generated by the three data comparisons identified a consistent set of adaptive pathways that contributed to the plastic traits involved in adaptation.

Earthworm genome methylation was found to be variable, suggesting a degree of stochasticity with functional plasticity. The identified miRNA networks showed a much higher relative earthworm site origin effect profile, relative to the transplant destination effect than did gene expression, suggesting expression sculpting via repression networks may act as a persistent functional memory within individuals exposed to environmental extremes. Among functional pathways, epithelial remodelling was identified as a physiological response and also independently as a functional signature in multiple omics datasets. This finding confirms this change as an acclimation response of *A. gracilis* to volcanic degassing soils. Circulatory system morphogenesis and angiogenesis were also repeatedly identified as functional acclimatisation responses, as was neuron development. Genes linked to these adaptive pathways were epigenetically modified to a far greater extent in earthworms experiencing environmental change (i.e. MV and VM treatments), regardless of the type of change. Signal transduction overall exhibited both a strong methylation response and gene expression profile changes. Metal accumulation was increased in degassing soils, and metal ion transporters were independently functionally identified as an acclimatisation response to degassing conditions.

Future studies should integrate comparative genomics of metal-detoxification gene families across earthworm species and experimentally quantify their inducible expression and methylation dynamics under controlled metal challenges, thereby resolving the relative contributions of metal tolerance and hypoxia acclimation to extreme-soil colonisation.

As critical ecosystem-service providers (Fonte, et al. 2023), annelids play a central role in soil formation, nutrient cycling, and carbon sequestration. By elucidating the molecular mechanisms that may underpin adaptive plasticity in *A. gracilis*, a framework is established for leveraging growing genomic and epigenomic resources to predict which species are likely to withstand or succumb to environmental extremes. This enhanced understanding will facilitate the development of molecular biomarkers for resilience or vulnerability, guide conservation priorities, and inform bioindicator design for monitoring ecosystem health under accelerating climate change.

## Acknowledgements

This work was funded by UKRI’s Natural Environment Research Council (GrantLNE/I026022/1). O.R. was supported by a UKRI GW4+ DTP studentship (NE/L002434/1). M.N. received a Marie Curie Fellowship (FP7-IEF-GA-2012-329690). L.C. was supported by a HorizonL2020 Marie SkłodowskalzlCurie Individual Fellowship (MSCAlzlIFlzl2014lzlGFlzl660378) and is currently funded by the Portuguese Foundation for Science and Technology (FCT; CEECIND/01986/2017). D.S. was funded through an IMP award from the UK Centre for Ecology & Hydrology Institutional Fund (ProjectL09221). We note with deep sadness that Professors Mike Bruford and Andrew John Morgan passed away during the preparation of this manuscript; their substantial contributions and enduring legacy remain integral to this work. This study was also supported by the strategic plan of the Centre for Functional Ecology, Science for People and the Planet (CFE) with the references UIDB/04004/2025 and UIDP/04004/2025, and by the strategic plan of the Associate Laboratory TERRA [LA/P/0092/2020] (https://doi.org/10.54499/LA/P/0092/2020).

## Data availability

All data used in this study have been deposited in the NCBI database under BioProject accession number PRJNA513445.

## Supplementary figure captions

Supplementary Fig. S1. Differential expression test results, effect-size shrunk log_2_ FC against normalised sample means for the RNAseq results.

Supplementary Fig. S2. Differential expression test results, effect-size shrunk log_2_ FC against normalised sample means for the miRNAseq normalised read count.

Supplementary Fig. S3. Differential expression test results, effect-size shrunk log_2_ FC against normalised sample means for the MeDIP-Seq results.

Supplementary Fig. S4. Rapid and bidirectional epidermal remodelling and mass change in *Amynthas gracilis* after reciprocal exposure to volcanic and reference soils. **a)** Individual body mass before and after 31 days of mesocosm exposure across four treatment groups: degassing-native maintained in degassing soil (V→V), degassing-native transplanted to reference soil (V→M), reference-native transplanted to degassing soil (M→V), and reference-native maintained in reference soil (M→M). Transplants from degassing to reference soil (V→M) gained weight, while those moved to degassing soils (M→V) showed marked weight loss, reflecting physiological stress. **b)** Representative epidermal histology in degassing-native worms (V) before and after exposure to reference soil (V→M). Epidermis thickened markedly after relocation to less stressful conditions. **c)** Quantification of epidermis thickness across treatment groups at the end of the 31-day experiment. Earthworms exposed to degassing soil (V→V, M→V) had significantly thinner epidermis compared to those in reference soil (V→M, M→M), regardless of origin. **d)** Representative histology of reference-native worms (M) before and after 31 days of exposure to volcanic soils (M→V), showing epidermal thinning after exposure to extreme conditions. Black scale bars in panels **b** and **d** = 25Lµm. Data support epidermal thickness as a plastic and rapidly adjusting morphological trait reflecting environmental stress, with coordinated changes in body mass.

Supplementary Fig. S5. Origin vs Origin differential miRNA expression network and functional enrichment clusters shows miRNAs (large dots) and genes (small dots) and whether or not they are upregulated (blue) or down-regulated (pink). All miRNA shown had a p<0.05 significance to its differential expression after fold-change effect size shrinking in deseq2.

Supplementary Fig. S6. Destination vs Destination differential miRNA expression network and functional enrichment clusters. Shows miRNAs (large dots) and genes (small dots) and whether they are upregulated (blue) or down-regulated (pink). All miRNA shown had a p<0.05 significance to its differential expression after fold-change effect size shrinking in deseq2.

---

**Supplementary Table S1.**
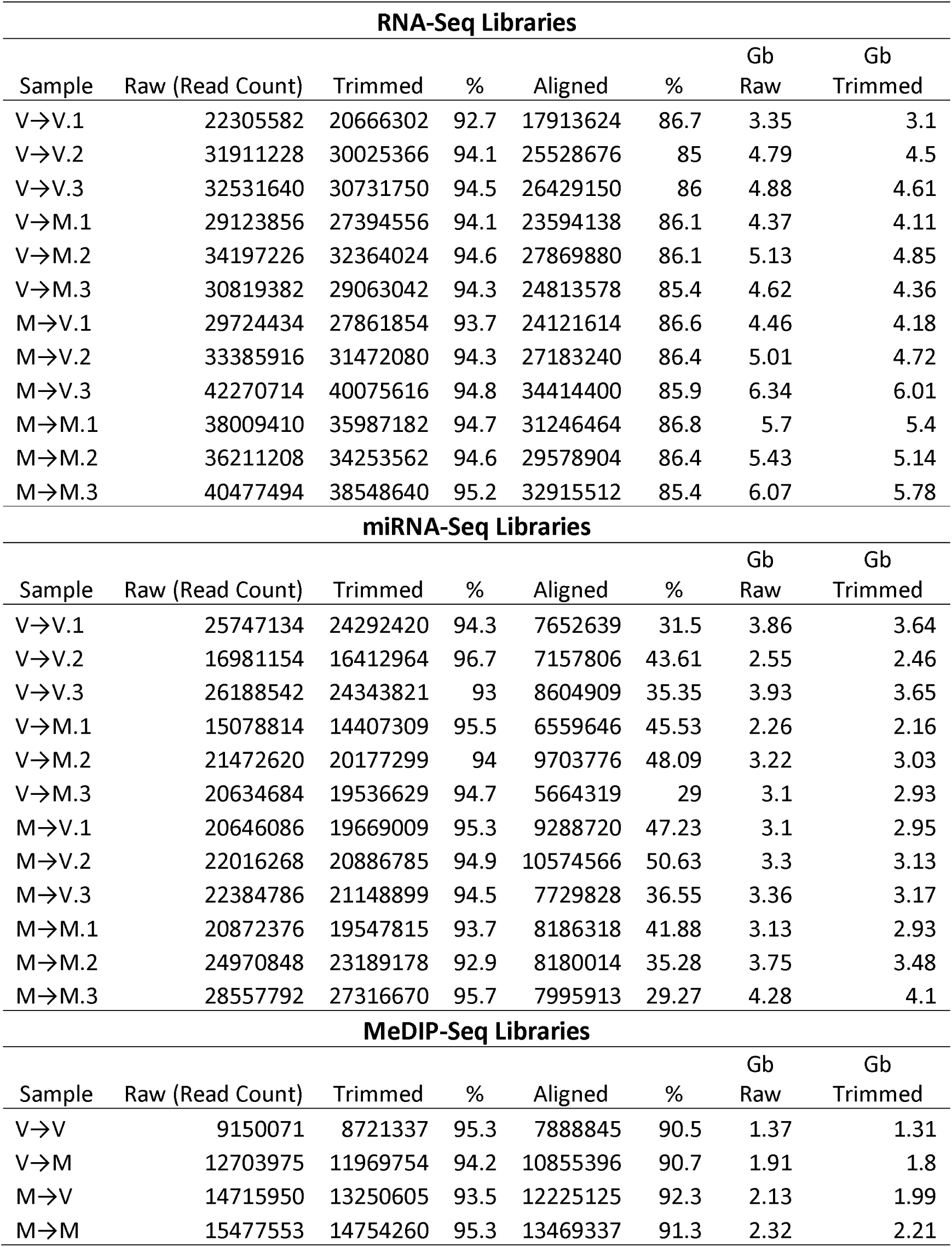
Sample sequencing libraries with trimming and alignment rates for the RNAseq, miRNA-seq and MeDIP-Seq libraries.

**Supplementary Table S2.** Gas and temperature conditions at the earthworm sampling and transplant deployment sites.

|  | Macela (Non-degassing site) |  |  |  |  |  | Furnas (Degassing site) |  |  |  |  |  |
| --- | --- | --- | --- | --- | --- | --- | --- | --- | --- | --- | --- | --- |
|  | Transplant Site |  |  | Collection Site |  |  | Transplant Site |  |  | Collection Site |  |  |
| Soil variable | Mean | Max | Min | Mean | Max | Min | Mean | Max | Min | Mean | Max | Min |
| CO<sub>2</sub> (vol%) 25 cm | nd | nd | nd | nd | nd | nd | 21.5 | 71.7 | 4.3 | 6.7 | 9.6 | 1.8 |
| CO<sub>2</sub> (vol%) 50 cm | 0.8 | 2.8 | 0.1 | 0.7 | 2.3 | 0.2 | 48.6 | 96.5 | 14 | 6.9 | 8.8 | 2.4 |
| CO<sub>2</sub> flow (g m<sup>-2</sup> d<sup>-1</sup>) | 19.5 | 27.5 | 12.3 | 15.4 | 32.4 | 6.2 | 181 | 533 | 58 | 19.8 | 25 | 12.5 |
| O<sub>2</sub> (vol%) 25 cm | 18.1 | 18.2 | 17.9 | - | - | - | 15.7 | 19.1 | 6.3 | 10.7 | 18.8 | 3.4 |
| O<sub>2</sub> (vol%) 50 cm | 18.5 | 19.9 | 13.1 | 18.8 | 20 | 15.6 | 10.4 | 17.1 | 1.4 | 9.9 | 18.3 | 1.2 |
| Temperature °C 25 cm | 17.5 | 17.6 | 16.2 | 17 | 18.9 | 15.9 | 35 | 45.7 | 28 | 18.4 | 19.6 | 17.9 |
| Temperature °C 50 cm | 17 | 17.6 | 16.2 | 17.1 | 18.8 | 15.9 | 37.3 | 47.3 | 29.8 | 17.8 | 19.3 | 17.8 |

**Supplementary Table S3.** Soil properties in soil samples collected from mesocosms. * Sample loss.

| Sample | Clay (%) | Silt (%) | Sand (%) | pH | Soil moisture content (%) | Loss on ignition (%) | Water holding capacity (%) |
| --- | --- | --- | --- | --- | --- | --- | --- |
| V→V.1 | 3 | 65 | 35 | 5.46 | 4.13 | 10.74 | 74.19 |
| V→V.2 | 2 | 66 | 35 | 5.69 | 7.47 | 10.71 | 78.93 |
| V→V.3 | 3 | 71 | 30 | 5.24 | 9.08 | 12.58 | 82.93 |
| V→V.4 | 5 | 70 | 29 | 5.4 | 5.2 | 12.64 | 93.5 |
| V→V.5 | 2 | 38 | 62 | 5.49 | 5.04 | 11.45 | 79.01 |
| V→V.6 | 3 | 66 | 35 | 5.83 | 4.83 | 10.25 | 75.43 |
| V→V Avg. | 3 | 62.67 | 37.67 | 5.52 | 5.96 | 11.4 | 80.67 |
| M→V.1 | 3 | 74 | 26 | 5.66 | 6.1 | 11.42 | 74.61 |
| M→V.2 | 4 | 76 | 25 | 5.43 | 5.51 | 11.11 | 86.15 |
| M→V.3 | 2 | 36 | 64 | 5.23 | 10.29 | 12.12 | 83.38 |
| M→V.4 | 3 | 68 | 33 | 5.84 | 8.08 | 10.87 | 85.42 |
| M→V.5 | 2 | 55 | 47 | 5.46 | 4.8 | 10.22 | 75.81 |
| M→V.6 | 2 | 62 | 40 | 5.53 | 5.32 | 9.89 | 76.81 |
| M→V Avg. | 2.67 | 61.83 | 39.17 | 5.53 | 6.68 | 10.94 | 80.36 |
| V→M.1 | 1 | 53 | 49 | 5.72 | 17.44 | 32.72 | 140.79 |
| V→M.2 | 1 | 54 | 47 | 5.6 | 8.17 | 24.52 | 106.36 |
| V→M.3 | 1 | 39 | 62 | 5.89 | 6 | 25 | 112.2 |
| V→M.4 | 2 | 50 | 51 | 5.45 | 13.39 | 29.37 | 120.16 |
| V→M.5 | 2 | 57 | 44 | 5.05 | 31.68 | 34.91 | 119.9 |
| V→M.6 | 1 | 62 | 40 | 5.65 | 16.48 | 23.06 | 106.4 |
| V→M Avg. | 1.33 | 52.5 | 48.83 | 5.56 | 15.53 | 28.26 | 117.63 |
| M→M.1 | 1 | 55 | 47 | 4.99 | 22.04 | 30.4 | 157.3 |
| M→M.2 | 1 | 50 | 51 | 5.23 | 12.92 | 25.61 | 103.99 |
| M→M.3 | 2 | 54 | 47 | 5.81 | 12.34 | 25.34 | 108.61 |
| M→M.4 | 1 | 53 | 49 | 5.33 | 22.56 | 36.7 | 155.31 |
| M→M.5 | 1 | 59 | 42 | 6 | 18.38 | 21.71 | 110.57 |
| M→M.6 | * | * | * | 6.37 | 5.8 | 14.42 | 100.16 |
| M→M Avg. | 1.2 | 54.2 | 47.2 | 5.62 | 15.67 | 25.7 | 122.66 |

## References

Alakarppa E, Salo HM, Valledor L, Cañal MJ, Häggman H, Vuosku J. 2018. Natural variation of DNA methylation and gene expression may determine local adaptations of Scots pine populations. Journal of Experimental Botany 69:5293–5305.

Andersen AC, Flores JF, Hourdez S. 2006. Comparative branchial plume biometry between two extreme ecotypes of the hydrothermal vent tubeworm Ridgeia piscesae. Canadian Journal of Zoology-Revue Canadienne De Zoologie 84:1810–1822.

Anderson C, Cunha L, Sechi P, Kille P, Spurgeon D. 2017. Genetic variation in populations of the earthworm, Lumbricus rubellus, across contaminated mine sites. BMC Genetics 18:97.

Arnold BE, Hodson ME, Charnock J, Peijnenburg W. 2008. Comparison of subcellular partitioning, distribution, and internal speciation of Cu between Cu-tolerant and naive populations of Dendrodrilus rubidus Savigny. Environmental Science & Technology 42:3900–3905.

Arnold R, Hodson M. 2007. Effect of time and mode of depuration on tissue copper concentrations of the earthworms Eisenia andrei, Lumbricus rubellus and Lumbricus terrestris. Environmental Pollution 148:21–30.

Bagnato E, Viveiros F, Pacheco J, D’agostino F, Silva C, Zanon V. 2018. Hg and CO2 emissions from soil diffuse degassing and fumaroles at Furnas Volcano (São Miguel Island, Azores): Gas flux and thermal energy output. Journal of Geochemical Exploration 190:39–57.

Banchroft J, Stevens A, Turner D. 1996. Theory and practice of histological techniques. 4th Edn New York. London, San Francisco, Tokyo: Churchil Livingstone.

Bartel DP. 2018. Metazoan MicroRNAs. Cell 173:20–51.

Baubron JC, Mathieu R, Miele G. 1991. Measurement of gas flows from soils in volcanic areas: the accumulation method. . International Conference on Active Volcanoes and Risk Mitig; Napoli.

Bhambri A, Dhaunta N, Patel SS, Hardikar M, Bhatt A, Srikakulam N, Shridhar S, Vellarikkal S, Pandey R, Jayarajan R, et al. 2018. Large scale changes in the transcriptome of Eisenia fetida during regeneration. Plos One 13.

Bolger AM, Lohse M, Usadel B. 2014. Trimmomatic: a flexible trimmer for Illumina sequence data. Bioinformatics 30:2114–2120.

Briones M, Álvarez-Otero R. 2018. Body wall thickness as a potential functional trait for assigning earthworm species to ecological categories. Pedobiologia 67:26–34.

Bushnell B. 2014. BBMap: A Fast, Accurate, Splice-Aware Aligner. 9th Annual Genomics of Energy & Environment Meeting.; March 17-20; United States.

Cai Y, Yu X, Hu S, Yu J. 2009. A brief review on the mechanisms of miRNA regulation. Genomics Proteomics Bioinformatics 147:147–154.

Camacho C, Coulouris G, Avagyan V, Ma N, Papadopoulos J, Bealer K, Madden TL. 2009. BLAST plus : architecture and applications. Bmc Bioinformatics 10.

Chang AJ, Bargmann CI. 2008. Hypoxia and the HIF-1 transcriptional pathway reorganize a neuronal circuit for oxygen-dependent behavior in Caenorhabditis elegans. Proceedings of the National Academy of Sciences of the United States of America 105:7321–7326.

Chiodini G, Cioni R, Guidi M, Raco B, Marini L. 1998. Soil CO2 flux measurements in volcanic and geothermal areas. Applied Geochemistry 13:543–552.

Cunha L, Campos I, Montiel R, Rodrigues A, Morgan AJ. 2011. Morphometry of the epidermis of an invasive megascoelecid earthworm (Amynthas gracilis, Kinberg 1867) inhabiting actively volcanic soils in the Azores archipelago. Ecotoxicology and Environmental Safety 74:25–32.

Cunha L, Montiel R, Novo M, Orozco-terWengel P, Rodrigues A, Morgan AJ, Kille P. 2014. Living on a volcano’s edge: genetic isolation of an extremophile terrestrial metazoan. Heredity 112:132–142.

Daniel O, Kohli L, Bieri M. 1996. Weight gain and weight loss of the earthworm Lumbricus terrestris L. at different temperatures and body weights. Soil Biology and Biochemistry 28:1235–1240.

Du Q, Luu P-L, Stirzaker C, Clark SJ. 2015. Methyl-CpG-binding domain proteins: readers of the epigenome. Epigenomics 7:1051–1073.

Edwards CA, Bohlen PJ. 1996. The Biology and Ecology of Earthworms. London, UK: Chapman and Hall.

Feinberg AP, Irizarry RA. 2010. Stochastic epigenetic variation as a driving force of development, evolutionary adaptation, and disease. Proceedings of the National Academy of Sciences of the United States of America 107:1757–1764.

Ferreira T, Gaspar JL, Viveiros F, Marcos M, Faria C, Sousa F. 2005. Monitoring of fumarole discharge and CO2 soil degassing in the Azores: contribution to volcanic surveillance and public health risk assessment. Annals of Geophysics 48:787–796.

Folmer O, Black M, Hoeh W, Lutz R, Vrijenhoek R. 1994. DNA primers for amplification of mitochondrial cytochrome c oxidase subunit I from diverse metazoan invertebrates. Molecular Marine Biology and Biotechnology 3:294–299.

Fonte SJ, Hsieh M, Mueller ND. 2023. Earthworms contribute significantly to global food production. Nature Communications 14:5713.

Friedländer MR, Mackowiak SD, Li N, Chen W, Rajewsky N. 2012. miRDeep2 accurately identifies known and hundreds of novel microRNA genes in seven animal clades. Nucleic Acids Research 40:37–52.

Friedman RC, Farh KKH, Burge CB, Bartel DP. 2009. Most mammalian mRNAs are conserved targets of microRNAs. Genome Research 19:92–105.

Gaspar J, Queiroz G, Ferreira T, Medeiros A, Goulart C, Medeiros J. 2015. Earthquakes and volcanic eruptions in the Azores region: geodynamic implications from major historical events and instrumental seismicity. In: Guest JE, Duncan, A.M., Barriga, F.J.A.S., Chester, D.K., editor. Volcanic Geology of São Miguel Island (Azores Archipelago). London: Geological Society of London Memoir. p. 33–49.

Gatzmann F, Falckenhayn C, Gutekunst J, Hanna K, Raddatz G, Carneiro VC, Lyko F. 2018. The methylome of the marbled crayfish links gene body methylation to stable expression of poorly accessible genes. Epigenetics & Chromatin 11.

Geeleher P, Huang RS, Gamazon ER, Golden A, Seoighe C. 2012. The regulatory effect of miRNAs is a heritable genetic trait in humans. BMC Genomics 13.

Griffiths-Jones S, Grocock RJ, van Dongen S, Bateman A, Enright AJ. 2006. miRBase: microRNA sequences, targets and gene nomenclature. Nucleic Acids Research 34:D140–D144.

Griffiths-Jones S, Saini HK, van Dongen S, Enright AJ. 2008. miRBase: tools for microRNA genomics. Nucleic Acids Research 36:D154–D158.

Guynes K, Sarre LA, Carrillo-Baltodano AM, Davies BE, Xu L, Liang Y, Martín-Zamora FM, Hurd PJ, de Mendoza A, Martín-Durán JM. 2024. Annelid methylomes reveal ancestral developmental and aging- associated epigenetic erosion across Bilateria. Genome Biology 25.

Haas BJ, Papanicolaou A, Yassour M, Grabherr M, Blood PD, Bowden J, Couger MB, Eccles D, Li B, Lieber M, et al. 2013. De novo transcript sequence reconstruction from RNA-seq using the Trinity platform for reference generation and analysis. Nature Protocols 8:1494–1512.

Höckner M, Dallinger R, Stüerzenbaum SR. 2015. Metallothionein gene activation in the earthworm (Lumbricus rubellus). Biochemical and Biophysical Research Communications 460:537–542.

Holt C, Yandell M. 2011. MAKER2: an annotation pipeline and genome-database management tool for second-generation genome projects. Bmc Bioinformatics 12.

Huang CD, Shen ZQ, Yue SZ, Jia L, Wang RP, Wang K, Qiao YH. 2023. Genetic evidence behind the Cd resistance of wild Metaphire californica: The global RNA regulation rather than specific mutation of well-known gene. Environmental Pollution 336.

Huang DW, Sherman BT, Tan Q, Kir J, Liu D, Bryant D, Guo Y, Stephens R, Baseler MW, Lane HC, et al. 2007. DAVID Bioinformatics Resources: expanded annotation database and novel algorithms to better extract biology from large gene lists. Nucleic Acid Research 35:W169–175.

ISO IOfS. 2005. Soil quality – Determination of pH. In. Geneva: International Organization for Standardization.

Jeltsch A, Jurkowska RZ. 2014. New concepts in DNA methylation. Trends in Biochemical Sciences 39:310–318.

Johnston MR, Herrick BM. 2019. Cocoon heat tolerance of Pheretimoid earthworms Amynthas tokioensis and Amynthas agrestis. American Midland Naturalist 181:299–309.

Kajitani R, Toshimoto K, Noguchi H, Toyoda A, Ogura Y, Okuno M, Yabana M, Harada M, Nagayasu E, Maruyama H, et al. 2014. Efficient de novo assembly of highly heterozygous genomes from whole- genome shotgun short reads. Genome Research 24:1384–1395.

Keller TE, Han P, Yi SV. 2016. Evolutionary transition of promoter and gene body DNA methylation across invertebrate-vertebrate boundary. Molecular Biology and Evolution 33:1019–1028.

Kille P, Andre J, Anderson C, Ang HN, Bruford MW, Bundy JG, Donnelly R, Hodson ME, Juma G, Lahive E, et al. 2013. DNA sequence variation and methylation in an arsenic tolerant earthworm population. Soil Biology & Biochemistry 57:524–532.

Klok C, Zorn M, Koolhaas JE, Eijsackers HJP, van Gestel CAM. 2006. Does reproductive plasticity in Lumbricus rubellus improve the recovery of populations in frequently inundated river floodplains? Soil Biology & Biochemistry 38:611–618.

Klupczynska EA, Ratajczak E. 2021. Can forest trees cope with climate change?-Effects of DNA methylation on gene expression and adaptation to environmental change. International Journal of Molecular Sciences 22.

Kozomara A, Birgaoanu M, Griffiths-Jones S. 2019. miRBase: from microRNA sequences to function. Nucleic Acids Research 47:D155–D162.

Kumari T, Phogat D, Jakhar N, Shukla V. 2024. Effectiveness of copper oxychloride coated with iron nanoparticles against earthworms. Scientific reports 14:23150.

Langdon CJ, Piearce TG, Meharg AA, Semple KT. 2003. Inherited resistance to arsenate toxicity in two populations of Lumbricus rubellus. Environmental Toxicology and Chemistry 22:2344–2348.

Langmead B, Trapnell C, Pop M, Salzberg SL. 2009. Ultrafast and memory-efficient alignment of short DNA sequences to the human genome. Genome Biology 10.

Lewis BP, Burge CB, Bartel DP. 2005. Conserved seed pairing, often flanked by adenosines, indicates that thousands of human genes are microRNA targets. Cell 120:15–20.

Li H, Handsaker B, Wysoker A, Fennell T, Ruan J, Homer N, Marth G, Abecasis G, Durbin R, Genome Project Data P. 2009. The Sequence Alignment/Map format and SAMtools. Bioinformatics 25:2078–2079.

Liebers R, Rassoulzadegan M, Lyko F. 2014. Epigenetic regulation by heritable RNA. PLOS Genetics 10.

Lindquist S, Craig EA. 1988. The heat-shock proteins. Annual review of genetics 22:631–677.

Lockwood BL, Julick CR, Montooth KL. 2017. Maternal loading of a small heat shock protein increases embryo thermal tolerance in Drosophila melanogaster. Journal of Experimental Biology 220:4492–4501.

Lokk K, Modhukur V, Rajashekar B, Märtens K, Mägi R, Kolde R, Koltsina M, Nilsson TK, Vilo J, Salumets A, et al. 2014. DNA methylome profiling of human tissues identifies global and tissue- specific methylation patterns. Genome Biology 15.

Lopienska-Biernat E, Stryinski R, Dmitryjuk M, Wasilewska B. 2019. Infective larvae of Anisakis simplex (Nematoda) accumulate trehalose and glycogen in response to starvation and temperature stress. Biology Open 8.

Love MI, Huber W, Anders S. 2014. Moderated estimation of fold change and dispersion for RNA-seq data with DESeq2. Genome Biology 15.

Luo MM, Sun LM, Hu J. 2009. Neural detection of gases - carbon dioxide, oxygen - in vertebrates and invertebrates. Current Opinion in Neurobiology 19:354–361.

Maegawa S, Hinkal G, Kim HS, Shen LL, Zhang L, Zhang JX, Zhang NX, Liang SD, Donehower LA, Issa JPJ. 2010. Widespread and tissue specific age-related DNA methylation changes in mice. Genome Research 20:332–340.

Mandrioli M. 2007. A new synthesis in epigenetics: towards a unified function of DNA methylation from invertebrates to vertebrates. Cellular and Molecular Life Sciences 64:2522–2524.

Mannello F, Medda V, Atonti G. 2011. Hypoxia and neural stem cells: from invertebrates to brain cancer stem cells. International Journal of Developmental Biology 55:569–581.

Martindale JL, Holbrook NJ. 2002. Cellular response to oxidative stress: Signaling for suicide and survival. Journal of Cellular Physiology 192:1–15.

Mendes EG, Nonato EF. 1957. The respiratory metabolism of tropical earthworms II. Studies on the cutaneous respiration. Boletim da Faculdade de Filosofia, Ciências e Letras, Universidade de São Paulo. Zoologia 21:153–166.

Monahan-Earley R, Dvorak AM, Aird WC. 2013. Evolutionary origins of the blood vascular system and endothelium. Journal of Thrombosis and Haemostasis 11:46–66.

Morgan AJ, Kille P, Stürzenbaum SR. 2007. Microevolution and ecotoxicology of metals in invertebrates. Environmental Science & Technology 41:1085–1096.

Morgan JE, Morgan AJ. 1990. The distribution of cadmium, copper, lead, zinc and calcium in the tissues of the earthworm Lumbricus rubellus sampled from one uncontaminated and 4 polluted soils. Oecologia 84:559–566.

Newbold LK, Robinson A, Rasnaca I, Lahive E, Soon GH, Lapied E, Oughton D, Gashchak S, Beresford NA, Spurgeon DJ. 2019. Genetic, epigenetic and microbiome characterisation of an earthworm species (Octolasion lacteum) along a radiation exposure gradient at Chernobyl. Environmental Pollution 255.

Novo M, Cunha L, Maceda-Veiga A, Talavera JA, Hodson ME, Spurgeon D, Bruford MW, Morgan AJ, Kille P. 2015. Multiple introductions and environmental factors affecting the establishment of invasive species on a volcanic island. Soil Biology & Biochemistry 85:89–100.

Novo M, Lahive E, Diez-Ortiz M, Spurgeon DJ, Kille P. 2020. Toxicogenomics in a soil sentinel exposure to Zn nanoparticles and ions reveals the comparative role of toxicokinetic and toxicodynamic mechanisms. Environmental Science-Nano 7:1464–1480.

Perez R, Aron S. 2023. Protective role of trehalose in the Namib desert ant, Ocymyrmex robustior. Journal of Experimental Biology 226.

Perry I, Hernadi SB, Cunha L, Short S, Marchbank A, Spurgeon DJ, Orozco-terWengel P, Kille P. 2022. Molecular insights into high-altitude adaption and acclimatisation of Aporrectodea caliginosa. Life Science Alliance 5.

Reiber CL, McGaw IJ. 2009. A review of the open and closed circulatory systems: New terminology for complex invertebrate circulatory systems in light of current findings. International Journal of Zoology. 2009: 301284 |.

Rieger RM, Purschke G. 2005. The coelom and the origin of the annelid body plan. Hydrobiologia 535:127–137.

Rowell DL. 1994. Soil Science: Methods and Applications. Harlow, UK: Longman Scientific and Technical.

Sarda S, Zeng J, Hunt BG, Yi SV. 2012. The evolution of invertebrate gene body methylation. Molecular Biology and Evolution 29:1907–1916.

Schachtman DP, Goodger JQD. 2008. Chemical root to shoot signaling under drought. Trends in Plant Science 13:281–287.

Schoville SD, Barreto FS, Moy GW, Wolff A, Burton RS. 2012. Investigating the molecular basis of local adaptation to thermal stress: population differences in gene expression across the transcriptome of the copepod Tigriopus californicus. Bmc Evolutionary Biology 12.

Shukla S, Kavak E, Gregory M, Imashimizu M, Shutinoski B, Kashlev M, Oberdoerffer P, Sandberg R, Oberdoerffer S. 2011. CTCF-promoted RNA polymerase II pausing links DNA methylation to splicing. Nature 479:74–79.

Silva C, Viveiros F, Ferreira T, Gaspar JL, Allard P. 2015. Diffuse soil emanations of radon and hazard implications at Furnas Volcano, São Miguel Island (Azores). In: Gaspar JL, Guest JE, Duncan AM, Barriga FJAS, Chester DK, editors. Volcanic Geology of São Miguel Island (Azores Archipelago): Geological Society of London. p. 0.

Silverman-Gavrila LB, Lu TZ, Prashad RC, Nejatbakhsh N, Charlton MP, Feng ZP. 2009. Neural phosphoproteomics of a chronic hypoxia model-Lymnaea stagnalis. Neuroscience 161:621–634.

Simao FA, Waterhouse RM, Ioannidis P, Kriventseva EV, Zdobnov EM. 2015. BUSCO: assessing genome assembly and annotation completeness with single-copy orthologs. Bioinformatics 31:3210–3212.

Srut M, Drechsel V, Hockner M. 2017. Low levels of Cd induce persisting epigenetic modifications and acclimation mechanisms in the earthworm Lumbricus terrestris. Plos One 12.

Suzuki MM, Kerr ARW, De Sousa D, Bird A. 2007. CpG methylation is targeted to transcription units in an invertebrate genome. Genome Research 17:625–631.

Swart E, Martell E, Svendsen C, Spurgeon DJ. 2022. Soil ecotoxicology needs robust biomarkers: A meta-analysis approach to test the robustness of gene expression-based biomarkers for measuring chemical exposure effects in soil invertebrates. Environmental Toxicology and Chemistry 41:2124–2138.

Talavera JA, Cunha L, Arévalo JR, Talavera IP, Kille P, Novo M. 2020. Anthropogenic disturbance and environmental factors drive the diversity and distribution of earthworms in Sao Miguel Island (Azores, Portugal). Applied Soil Ecology 145.

Viljoen SA, Reinecke AJ. 1992. The temperature requirements of the epigeic earthworm species Eudrilus eugeniae (Oligochaeta) - a laboratory study. Soil Biology and Biochemistry 24:1345–1350.

Viveiros F, Baldoni E, Massaro S, Stocchi M, Costa A, Caliro S, Chiodini G, Andrade C. 2023. Quantification of CO2 degassing and atmospheric dispersion at Caldeiras da Ribeira Grande (São Miguel Island, Azores). Journal of Volcanology and Geothermal Research 438:107807.

Viveiros F, Cardellini C, Ferreira T, Caliro S, Chiodini G, Silva C. 2010. Soil CO2 emissions at Furnas volcano, São Miguel Island, Azores archipelago: Volcano monitoring perspectives, geomorphologic studies, and land use planning application. Journal of Geophysical Research: Solid Earth 115.

Viveiros F, Ferreira T, Vieira JC, Silva C, Gaspar JL. 2008. Environmental influences on soil CO_2_ degassing at Furnas and Fogo volcanoes (Sao Miguel Island, Azores archipelago). Journal of Volcanology and Geothermal Research 177:883–893.

Viveiros F, Gaspar JL, Ferreira T, Silva C, Marcos M, Hipólito A. 2015. Mapping of soil CO2 diffuse degassing at São Miguel Island and its public health implications. In: Gaspar JL, Guest JE, Duncan AM, Barriga FJAS, Chester DK, editors. Volcanic Geology of São Miguel Island (Azores Archipelago): Geological Society of London. p. 0.

Wang C, Sun Z, Zheng D, Liu X. 2011. Function of mucilaginous secretions in the antibacterial immunity system of Eisenia fetida. Pedobiologia 54:S57–S62.

Wang K, Qiao YH, Li HF, Huang CD. 2020. Use of integrated biomarker response for studying the resistance strategy of the earthworm Metaphire californica in Cd-contaminated field soils in Hunan Province, South China. Environmental Pollution 260.

Wang X, Li A, Wang W, Que H, Zhang G, Li L. 2021. DNA methylation mediates differentiation in thermal responses of Pacific oyster (Crassostrea gigas) derived from different tidal levels. Heredity 126:10–22.

Wilson GA, Beck S. 2016. Computational Analysis and Integration of MeDIP-seq Methylome Data. In. Next Generation Sequencing: Advances, Applications and Challenges.: InTechOpen. p. 153–169.

Zhao X, Yu H, Kong L, Liu S, Li Q. 2016. High throughput sequencing of small RNAs transcriptomes in two Crassostrea oysters identifies microRNAs involved in osmotic stress response. Scientific reports 6:22687.

