## Supplementary figures and images for "Coping with extremes: How Epigenetic and Molecular Adaptations Enable Earthworms to Thrive in Volcanic Soils"

### Supplementary Fig S1

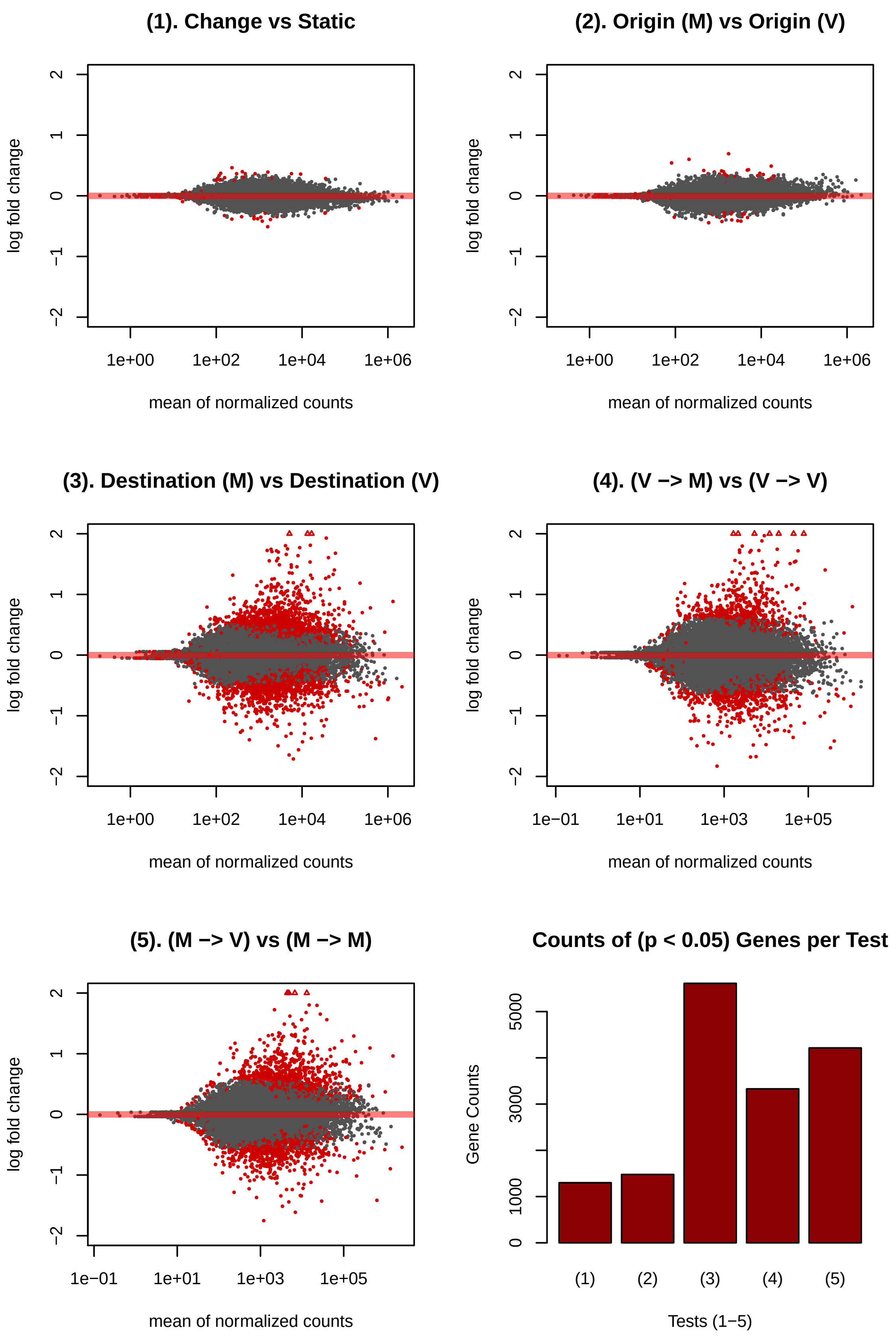

### Supplementary Fig S2

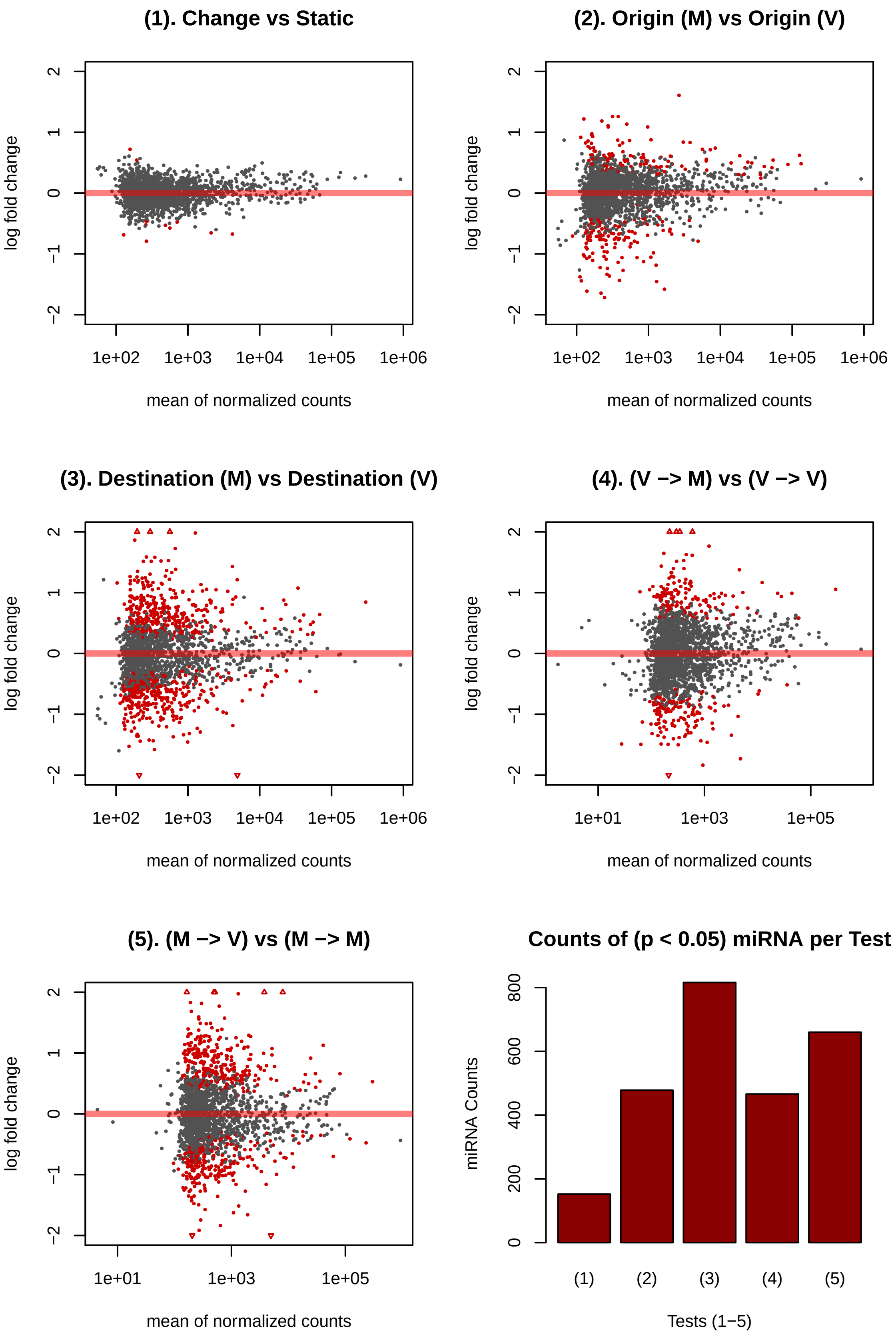

### Supplementary Fig S3

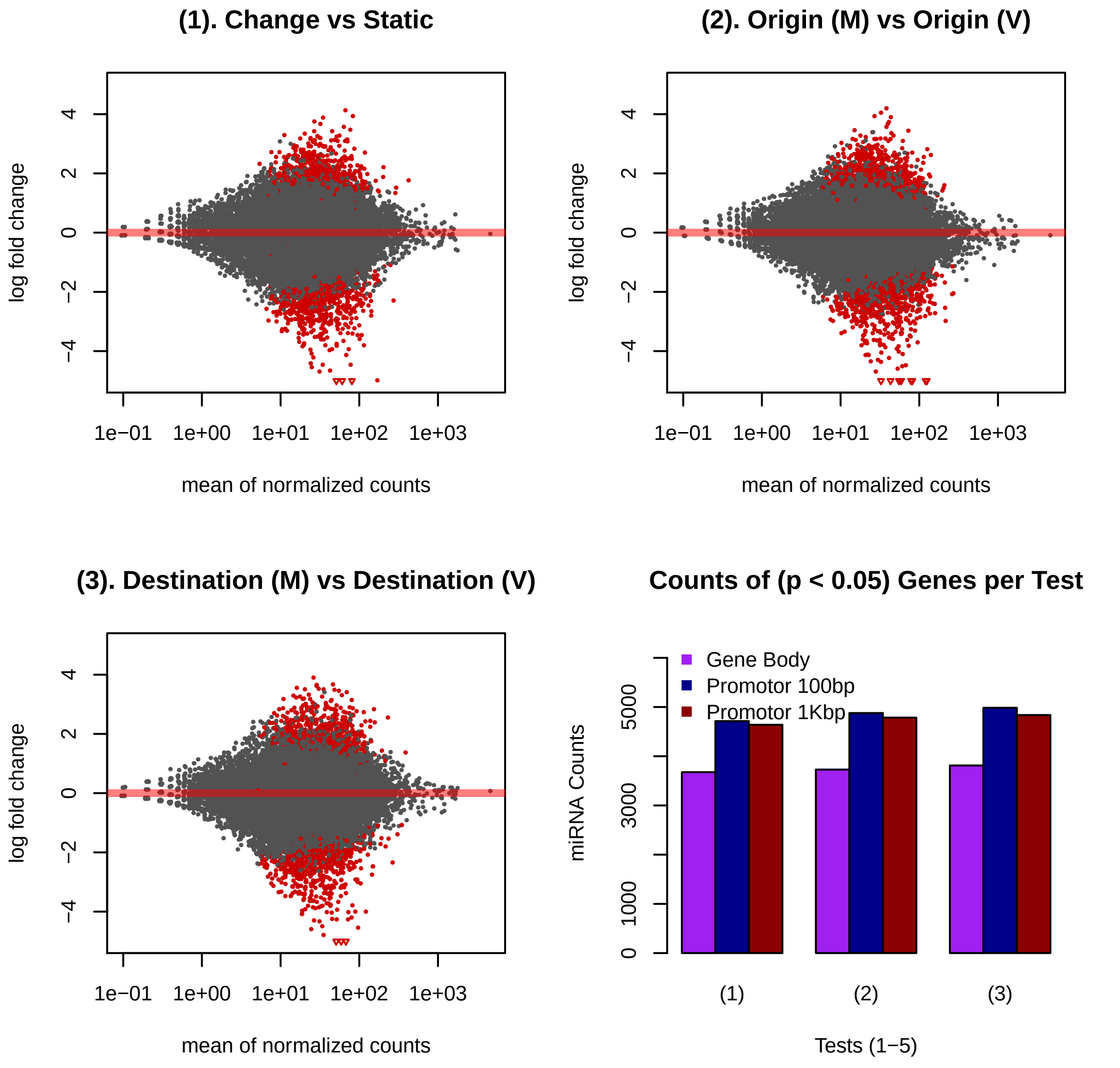

### Supplementary Fig S4

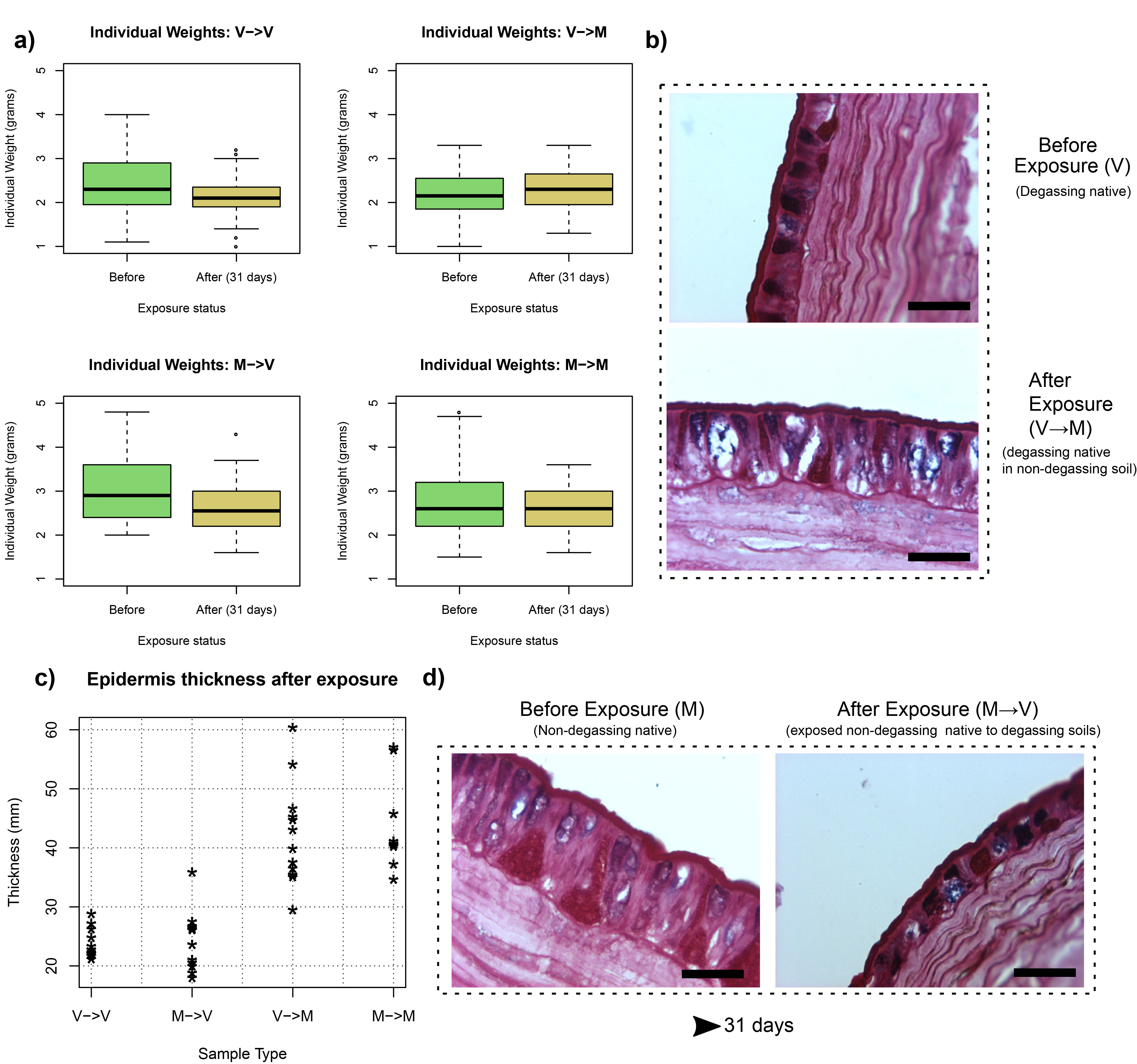

### Supplementary Fig S5

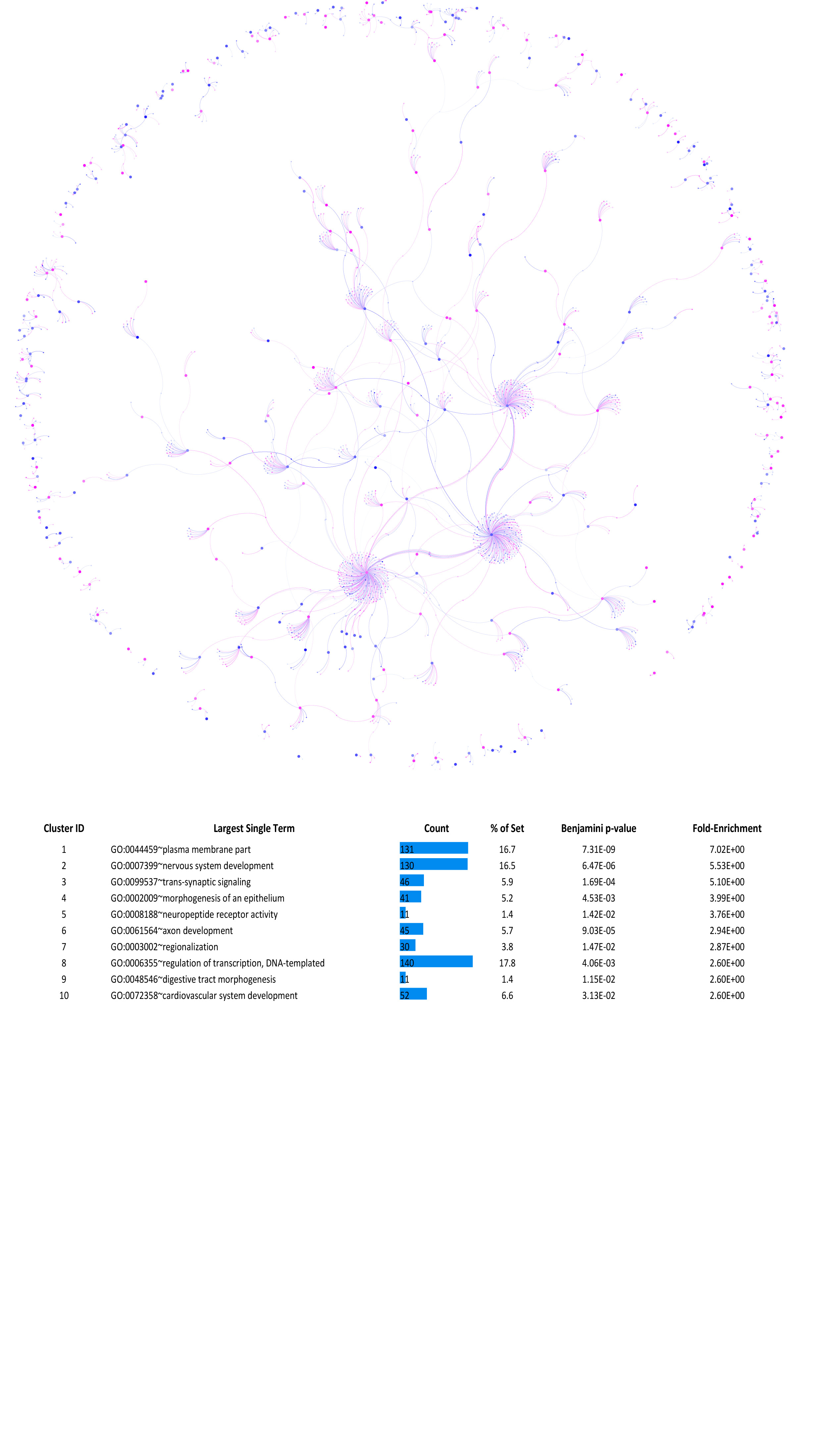

### Supplementary Fig S6

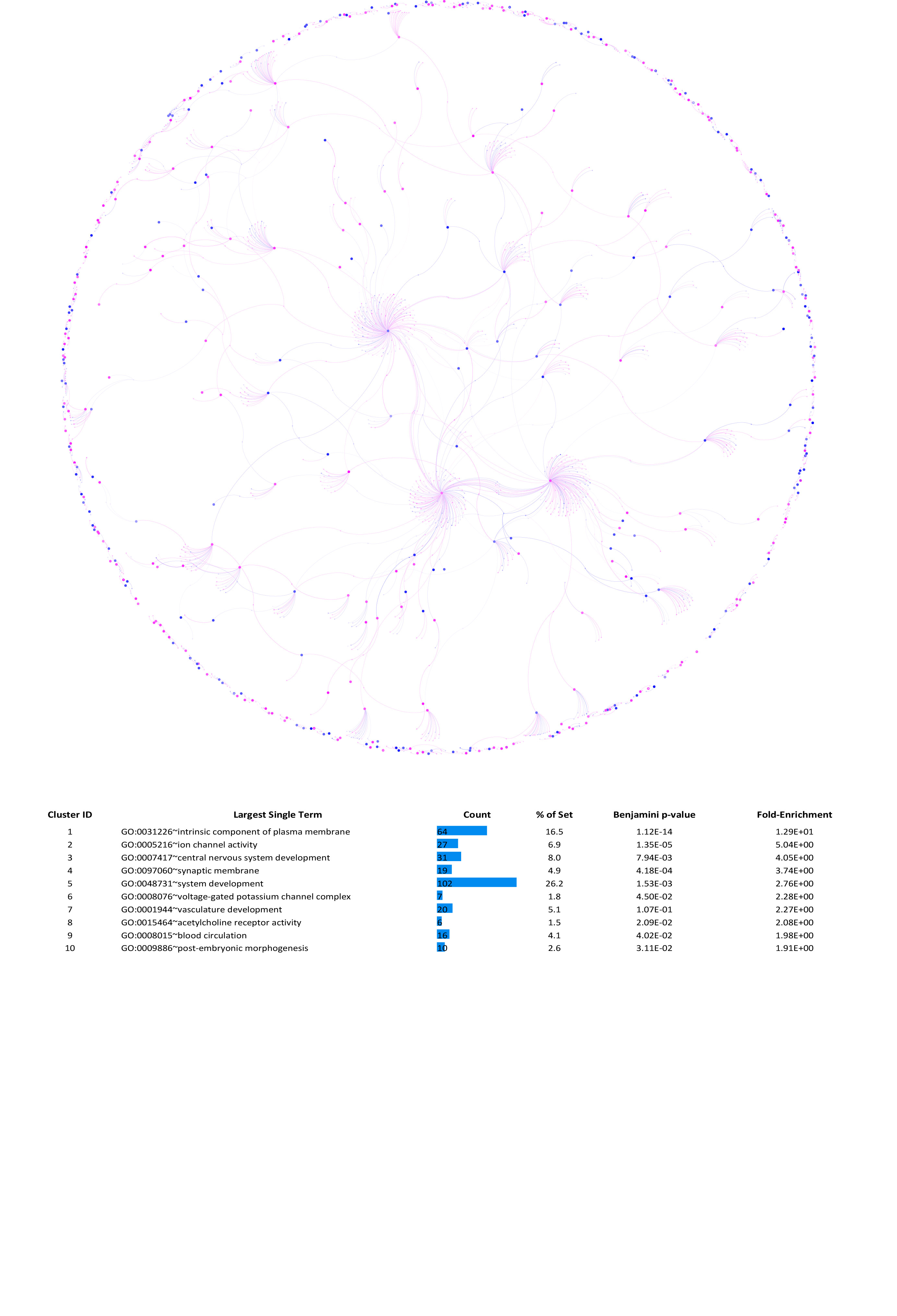
